# Spatiotemporal distribution of neural crest cells in the common wall lizard *Podarcis muralis*

**DOI:** 10.1101/2024.05.24.595691

**Authors:** Robin Pranter, Nathalie Feiner

## Abstract

**Background:** Neural crest cells (NCCs) are migratory embryonic stem cells that give rise to a diverse set of cell types. Here we describe the dynamic distribution of NCCs in developing embryos of the common wall lizard *Podarcis muralis* inferred from ten markers. Our aim is to provide insights into the NCC development of lacertid lizards and to infer evolutionary modifications by comparisons to other tetrapods.

**Results:** NCC migration is ongoing at oviposition, following three streams in the head and multiple in the trunk. From 21ss, we observe expression patterns indicating the beginning of differentiation towards mesenchymal and neuronal fates. By 35ss, migration is restricted to caudal levels, and fully differentiated chromaffin cells are observed.

**Conclusions:** We find that some markers show patterns that differ from other tetrapods. For example, the antibody HNK-1 labels three NCC streams from the hindbrain while some comparable reptile studies describe four. However, the information emerging from all markers combined shows that the overall spatiotemporal distribution of NCCs in the common wall lizard is largely conserved with that of other tetrapods. Our study highlights the dynamic nature of seemingly canonical marker genes and provides the first description of spatiotemporal NCC dynamics in a lacertid lizard.

## Introduction

Neural crest cells (NCCs) are transient, migratory stem cells that are unique to vertebrates and that give rise to many different cell types, thus contributing to a diversity of traits. NCCs are specified in the dorsal neural tube from where they delaminate in an epithelial-to-mesenchymal transition (EMT) and subsequently migrate throughout the embryo and differentiate. The cell types they give rise to include chromatophores responsible for skin color, osteocytes and chondrocytes that constitute the facial skeleton, neurons and glia of the peripheral nervous system and endocrinal chromaffin cells in the adrenal gland.^1^

NCCs originated at the dawn of vertebrates, and many of the vertebrate key innovations (most notably the jaw and other craniofacial structures) develop from NCCs.^2^ For this reason, a substantial body of research has targeted the evolutionary origin of NCCs by investigating NCCs in early branching vertebrates and NCC-like cells in non-vertebrate chordates.^3–5^ In addition to these attempts to elucidate the evolutionary origin of NCCs in a macro-evolutionary context, much research has focused on investigating their function and development in a number of model organisms, most notably mouse (*Mus musculus*), chicken (*Gallus gallus*), African clawed frog (*Xenopus laevis*) and zebrafish (*Danio rerio*). Motivation for this research partially stems from the fact that failure of NCC functions cause neurocristopathies, a highly heterogeneous family of congenital diseases,^6^ which makes them relevant from a medical perspective.

In squamate reptiles (lizards and snakes), NCC biology has been investigated in three species: the California kingsnake (*Lampropeltis getula californiae*^7^), the Egyptian cobra (*Naja haje haje*^8^) and the veiled chameleon (*Chamaeleo calyptratus*^9^). The reports on the two snake species mainly focus on the migration of NCCs in the trunk, while the report in chameleon provides a comprehensive description of both cranial and trunk NCCs. Both snakes and chameleons have highly derived body plans including in NCC-derived traits such as skull morphology.

In the chameleon, cranial NCC migration starts between somite stage 4 and 6 (4ss and 6ss),^9^ which is similar to mouse (5ss^10–12^), human (4ss^13^) and chicken (6ss^14(p66)^). In general, cranial NCC migration typically follows three main dorsolateral streams.^15^ The NCCs in the anterior-most stream are specified in the caudal midbrain and the first three rhombomeres of the hindbrain and contribute cells to the rostrum, maxillary process and first pharyngeal arch.^15^ The second stream is constituted by NCCs that are specified in rhombomeres three through five (anterior to the otic vesicle) and migrate to the second pharyngeal arch.^15^ And lastly, the NCCs in the third stream are specified in rhombomeres five and six (posterior to the otic vesicle) and migrate to the third pharyngeal arch.^15^ In the chameleon, however, Diaz et al.^9^ identified a fourth stream of cranial NCCs starting posteriorly to the third stream and migrating to the fourth pharyngeal arch. An equivalent stream has also been described in alligator (*Alligator mississippiensis*) and ostrich (*Struthio camelus*).^16^

NCC migration in the trunk follows multiple, periodic streams starting in the dorsal neural tube. Viewed in cross-sections in a transverse plane, these streams typically follow two different routes.^17–19^ An early wave of migrating cells follows a ventromedial route traversing between somites and through the anterior portion of somites and giving rise to, for example, peripheral nervous system and chromaffin tissue.^17–19^ A later wave follows a dorsolateral route and gives rise to chromatophores.^17–19^ While this pattern seems to be largely conserved across tetrapods,^7–9^ it has been reported that some NCCs in the second wave follow a ventromedial rather than dorsolateral route in the California kingsnake.^7^ The equivalent was not reported for chameleon or cobra.^8,9^

In between and partially overlapping with the cranial and trunk NCCs, there is a small subpopulation of NCCs called vagal NCCs.^15^ Similar to trunk NCCs, vagal NCCs migrate along both dorsolateral and ventromedial routes.^20^ The dorsolateral migrating NCCs reach the pharyngeal arches and the heart.^20^ Some vagal NCCs of both the dorsoventral and ventromedial streams reach the foregut which they then follow caudally.^20^ Vagal NCCs have not been described in squamates.

While efforts in squamates have increased our understanding of NCC specification, migration and differentiation across vertebrates, evolutionary changes in NCC behavior and potentially associated phenotypic changes are currently underexplored.^21^ Since NCCs give rise to functionally diverse and ecologically relevant traits ranging from coloration (e.g., pigment cells) and morphology (e.g., skull and jaw bones) to physiology and behavior (e.g., adrenal gland influencing aggression), their coupling in a shared developmental origin can potentially have evolutionary consequences. Indeed, this coupling has been suggested to explain the domestication syndrome, which is based on the observation that domesticated animals share a set of changes in NCC-derived traits.^22,23^ In general, due to their central role during development, modifications in the regulation of NCCs are expected to have the potential to generate biases in micro-evolutionary variation in natural populations of vertebrates and therefore contribute to adaptation and diversification.^21,24,25^ To evaluate hypotheses like this, establishing patterns of NCC migration in species where NCCs may be implied in micro-evolutionary adaptation is crucial.

Here we present a description of NCCs in the common wall lizard (*Podarcis muralis*), which is a small (approx. 50–70 mm snout-to-vent length^26^), egg-laying lacertid. This species belongs to a genus of lizards distributed in the Mediterranean and Southern/Central Europe that shows high variation in coloration and color patterns.^27,28^ Within the common wall lizard, there is substantial variation in predominantly NCC-derived traits, namely coloration (ranging from brown/tan to black/green),^29–32^ social behavior (in particular aggression in male-male interactions),^33,34^ and morphology (e.g., body size, head size and shape).^29,32^ In Italy, the strength of expression of these traits is tightly correlated across the landscape, culminating in the *nigriventris* syndrome at one end of the extreme and the ancestral phenotype at the other end.^32^ The genetic basis of these trait differences has been identified as being polygenic, with more than half of all identified genes having a known association with the regulation of NCCs.^32^ This prompted us to investigate NCC biology in this species. Females typically lay 2-3 clutches per breeding season with 2-10 eggs per clutch.^26(p189)^ We use a set of ten NCC markers that target different aspects of NCC biology to investigate the spatial distribution and infer migration patterns of NCCs from the earliest stage at oviposition (∼13ss) until the latest stages of NCC activity (∼ 64ss). To identify evolutionary changes to the spatiotemporal distribution of NCCs, we compare our observations to equivalent patterns described in other squamates and more distantly related tetrapods. We find broad conservation in general NCC specification, migration, and differentiation, but also taxonomic differences in the expression patterns of individual markers. Taken together, our description of spatial and temporal patterns of NCCs during embryonic development expands our understanding of the taxonomic diversity in NCC biology and presents a first step towards investigating the role of NCCs in shaping micro-evolutionary patterns.

## Results and Discussion

Given that NCCs are a heterogeneous and highly dynamic cell population, there is no single marker that distinguishes them from other cell types throughout development. We therefore use a two-pronged approach to resolve spatiotemporal dynamics of putative NCCs. Firstly, we use immunostaining with antibodies against Human Natural Killer 1 (HNK-1) to label putative NCCs throughout development. HNK-1 is a carbohydrate on the cell surface of NCCs that is involved in cell-cell communication during migration^35,36^ and it is a widely used marker of newly formed and migrating NCCs^16,17,37–43^ (note however that it is not labelling NCCs in mouse)^44^. This antibody has been successfully applied in other squamate species,^7–9^ which allows a straightforward comparison between our observations in the common wall lizard and other squamates.

Since HNK-1 is known to also label other cell types, in particular at late developmental stages,^45^ we complement HNK-1 immunostainings with gene expression analyses of NCC marker genes. For this second approach, we select a panel of nine marker genes whose expression patterns are visualized using *in situ* hybridizations. Collectively, these marker genes label the majority of NCC subpopulations and target various stages of NCC development.^46^ Note however that some markers do not exclusively label NCCs since they fulfil pleiotropic functions (e.g., *Twist1* stains mesodermal cells in addition to NCCs^47^). Specifically, we select *Wnt1* and *Snai2* to label predominantly early NCCs, *Snai1*, *Twist1,* and *Sox9* to mark NCCs biased towards mesenchymal fates and *FoxD3*, *Sox10 and Phox2b* to mark NCCs biased towards neuronal and glial fates.^46,48,49^ In addition, we use *Tyrosine hydroxylase* (*Th*) to label NCCs differentiating into chromaffin cells.^50^ While HNK-1 immunostaining and *in situ* hybridizations provide snapshots of the location of NCCs throughout development, we infer migratory routes from the series of observed imaging data aided by comparisons to relevant literature. For clarity, results for cranial, vagal (where applicable) and trunk NCCs are presented separately for each marker.

### HNK-1 stains migratory and early differentiating NCCs

HNK-1-positive cells are present in all investigated stages (five stages ranging from 13ss to 35ss; Figure 1A-P). Cranial NCC migration can be inferred along three routes. Each stream originates in the dorsal midline and extends ventrally in the developing head (Figure 1 A, B, D, E and J; dashed arrows indicate inferred streams). The first stream starts at the level of the midbrain-hindbrain boundary and reaches towards the rostrum, maxillary process and the first pharyngeal arch (Figure 1A). The second and third inferred streams originate in the hindbrain on either side of the otic vesicle, extending towards the second and third pharyngeal arches respectively (Figure 1D and E). While the first stream is visible already at the first investigated stage (13ss), the second and third streams are seen more clearly from 17ss. The three streams of migrating cranial NCCs remain recognizable until 22ss (Figure 1S), after which migration of cranial NCCs presumably seizes. By then, HNK-1 signal is widely spread in the developing head including structure that will give rise to the facial skeleton such as the rostrum, maxillary process, and the first three pharyngeal arches (Figure 1J), and some cranial placodes (Figure 1N), which are co-derived from NCCs and placodal cells.^56,57^ HNK-1 signal in the cranial peripheral nervous system is maintained as the cranial placodes give rise to cranial nerves (Figure 1O; cranial nerve nomenclature follows Diaz et al. (2019),^9^ Lee et al. (2003)^57^ and Streeter (1905)^58^). There is also strong segmental staining in the rhombomeres of the hindbrain (Figure 1K and L).

**Figure 1.**
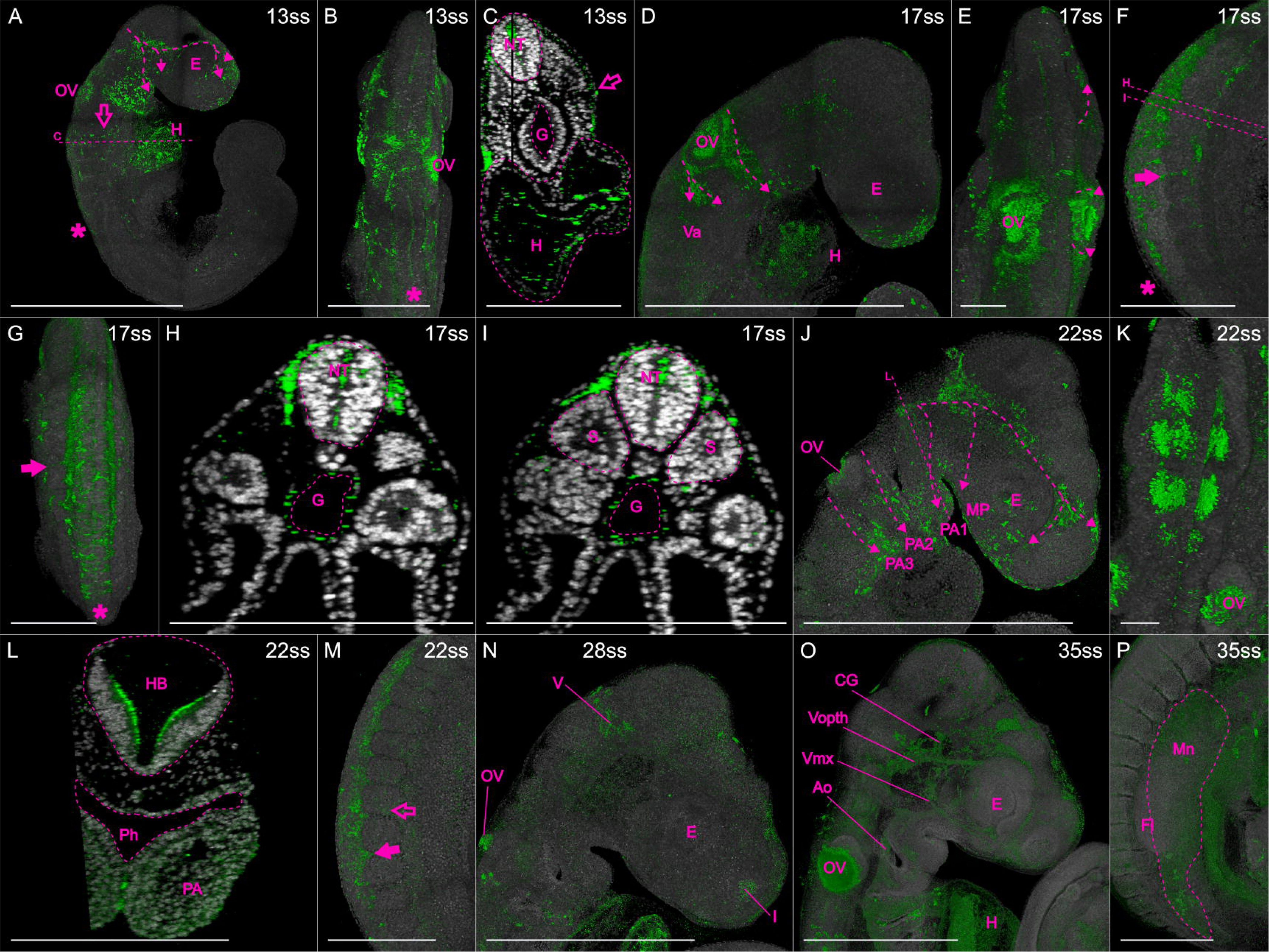
Immunohistochemistry of HNK-1 stains migratory and some early differentiating NCCs in common wall lizard embryos. Dashed arrows indicate inferred cranial NCC migration. **A-C** 13ss embryo in lateral (**A**) and dorsal (**B**) view, and in optical cross-section (**C**). Asterisks indicate expression in premigratory and early migratory NCCs, and open arrows indicate vagal NCCs. Staining in the heart is best seen in **A** and **C. D-E** Head of 17ss embryo in lateral (**D**) and dorsal (**E**) view. **F**-**I** Trunk of 17ss embryo in lateral (**F**) and dorsal (**G**) view, and in optical cross-sections (**H** and **I**). Arrows indicate inter-somitic NCC migration and asterisks indicate premigratory and early migratory NCCs. **J-L** Head of 22ss embryo in lateral (**J**) and dorsal (**K**) view, and in optical cross-section (**L**). **M** Trunk of 22ss embryo in lateral view. Arrow indicates inter-somitic NCC migration and open arrow indicates migration at the level of a somite. **N** and **O** Heads of 28 (**N**) and 35ss (**O**) embryos in lateral view. **P** Trunk of 35ss embryo in lateral view. **O** and **P** were rendered with lower DAPI-opacity. The locations of the optical cross sections **C**, **H**, **I** and **L** are indicated in **A**, **F** and **J**. **Abbreviations:** E, eye; Fb, forebrain; Fl, Forelimb; G, gut; H, heart; HB, hindbrain; Mn, mesonephros; MP, maxillary process; NT neural tube; OV, otic vesicle; PA, pharyngeal arch; Ph, pharynx; S, somite; Va, vagal. **Cranial nerves:** I, olfactory nerve (also known as cranial nerve 1); V, trigeminal nerve (also known as cranial nerve 5); Vmx, maxillary branch of the trigeminal nerve; Vopth, ophthalmic branch of the trigeminal nerve. **Scale bars** in **B**-**D**, **F**-**L** and **O**-**P** are 0.5 mm, all other scale bars are 1 mm.

At vagal levels, NCC migration can first be inferred by 13ss along a dorsolateral route to the heart (open arrows in Figure 1A and C), which is also strongly stained by HNK-1 (indicated with “H” in Figure 1A, C and D). At later developmental stages, there is a branch from the third cranial stream extending caudally into the trunk, which potentially indicates a stream of vagal NCCs on their way to populate the gut and give rise to the enteric nervous system (indicated with “Va” in Figure 1D). Consistently, cross-sections of the trunk reveal staining around the gut (Figure 1H-I); a similar staining has been shown in chicken.^38^

In the trunk, HNK-1 signal in NCCs can be roughly divided into four stages: premigratory, early migratory, migratory, and early neural differentiation. HNK-1 signal is present in a stripe along the dorsal midline where premigratory NCCs are expected (Figure 1B, E and G). Immediately lateral to the dorsal midline, HNK-1 stains early migratory NCCs in loose cell aggregates, not yet forming migratory streams (Figure 1A, B, F, G and M). The staining of premigratory and early migratory NCCs is found at progressively more posterior positions along the trunk as development proceeds. At more anterior levels, multiple streams of NCCs can be inferred between the somites (arrows in Figure 1F, G and M), and, to a lesser extent, at the level of the somites (open arrow in Figure 1M). Cross-sections reveal that the streams of migratory NCCs take a ventral route between somites (inter-somitic; Figure 1H), while they take a dorsolateral route at the levels of the somites (Figure 1I). NCC migration in the most caudal region of the trunk is still ongoing in the oldest investigated stages (28ss and 35ss). While HNK-1 signal is known to mark the NCC-derived dorsal root ganglia in chicken,^38^ and also a turtle (*Trachemys scripta*)^59^, an alligator (*Alligator mississippiensis*)^16^ and the Californian kingsnake (*Lampropeltis getula calilforniae*),^7^ this staining is absent in common wall lizards (Figure 1P). The reported HNK-1 staining in dorsal root ganglia in chameleon is also less distinct than in other organisms.^9^ This emphasizes that HNK-1 is neither specific, nor general enough to figure as a sole NCC marker, and therefore requires complementation with other markers. In later stages, there is some HNK-1 staining in the mesonephros (Figure P). This is also seen in other reptiles, but it is unclear if it is due to NCCs contributing to this organ.^7,9^ At later stages (35ss), we detect HNK-1 signal in blood cells inside the aorta and heart (Figure 1O). Presumably, this is caused by the fact that blood cells possess HNK-1 on their cell surface, despite not being NCC-derived.^45^

### *Wnt1* and *Snai2* are expressed in premigratory NCCs

Besides being a crucial organizer of the midbrain-hindbrain region,^60^ *Wnt1* is a classic marker of the neural border (including the rhombic lips^61^) and inducer of NCC specification.^46,62^ And the *Wnt1-Cre* transgenic lines have been instrumental in numerous experimental studies of NCCs in mice,^48,63,64^ highlighting its crucial role as a NCC regulator.

*Wnt1* is expressed in all investigated stages (10 stages ranging from 13ss to 60ss), mostly along the dorsal midline (Figure 2A-E). Through most of the investigated developmental stages (16-30ss), there is a noticeable gap of expression in the most anterior part of the hindbrain (the metencephalon; asterisks in Figure 2A-D), separating the expression into an anterior and a posterior domain. This pattern has been described in chicken.^60^ The gap of expression overlaps with the origin of the first cranial stream as inferred from HNK-1 staining.

**Figure 2.**
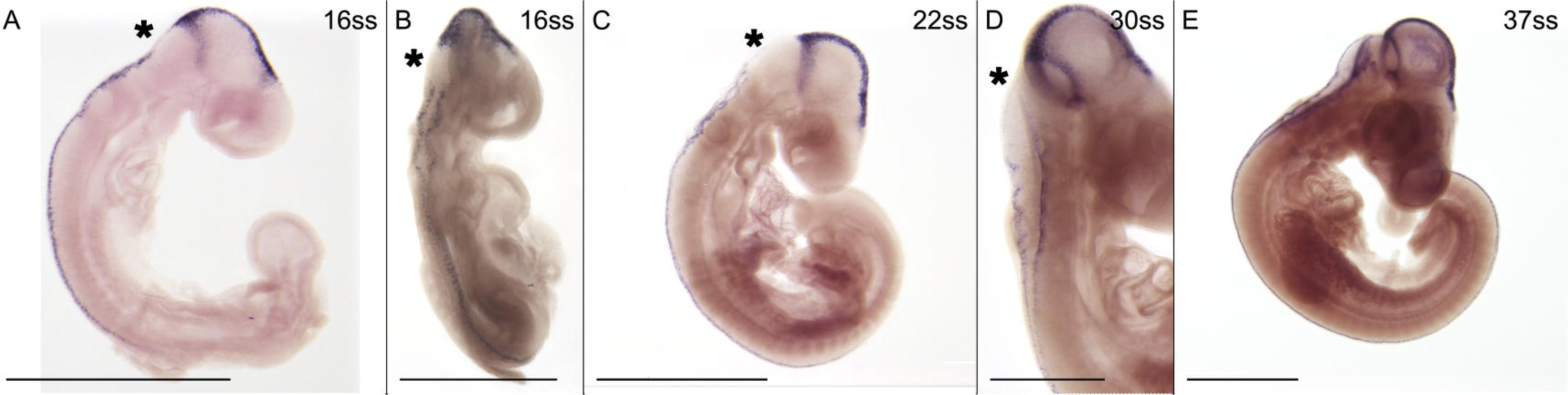
*Wnt1* is expressed along the dorsal midline. Asterisks indicate a gap with no expression in the anterior hindbrain. **A, B** 16ss embryo in lateral (**A**) and dorsolateral (**B**) view. **C** 22ss embryos in lateral view. **D** Hindbrain of 30ss embryo in dorsolateral view. **E** 30ss embryos in lateral view. **Scale bars** in **B** and **D** are 0.5 mm, all other scale bars are 1 mm.

The anterior domain runs along the dorsal midline of the head from the level of the diencephalon to the midbrain-hindbrain boundary (Figure 2A, C). It has some lateral extensions that are presumably not NCC related and will not be described in further detail. The posterior domain consists of two parallel lines running along the rhombic lips in the hindbrain (Figure 2D) and continuing caudally along the dorsal midline of the trunk (Figure 2B), extending caudally approximately as far as to the last formed somite pair, and continuously extends posteriorly as development progresses (Figure 2A, C and E). The caudal extension of *Wnt1* expression is concordant with premigratory NCCs as inferred from HNK-1 staining.

*Snai1* and *-2* are classical NCC markers previously known as *Snail* and *Slug*.^65^ They are transcription factors that control cadherin-transcription and thereby EMT.^46^ There was some uncertainty in regards to the homology relationships between these genes, and EMT is induced by paralogous genes in chicken (*Snai2*) and mouse (*Snai1*).^66^ We confirm the expected orthology relationships between *Snai1* and -*2* of the common wall lizard and their homologs in human, mouse and chicken (see Methods) and investigate both of them using *in situ* hybridization. *Snai2* expression is investigated in eight stages ranging from 13ss to 51ss.

*Snai2* expression in the head is found in putative premigratory NCCs along the dorsal midline and the rhombic lips in the hindbrain (Figure 3A and B), and in the three streams of cranial NCC migration previously inferred with HNK-1 staining (dashed arrows in Figure 3A and C). In cross-section through the second stream, *Snai2* expression can be seen from the dorsal tip of the neural crest to the pharyngeal arches (Figure 3C). While a caudally migrating vagal stream of NCCs is only weakly suggested by HNK-1, it is strongly marked by *Snai2* expression, visible as a branch from the third cranial stream (Figure 3A). In cross-section at the level of the heart, strong *Snai2* expression is located lateral to the gut, presumably indicating caudally migrating vagal NCCs (Figure 3D).^20^

**Figure 3.**
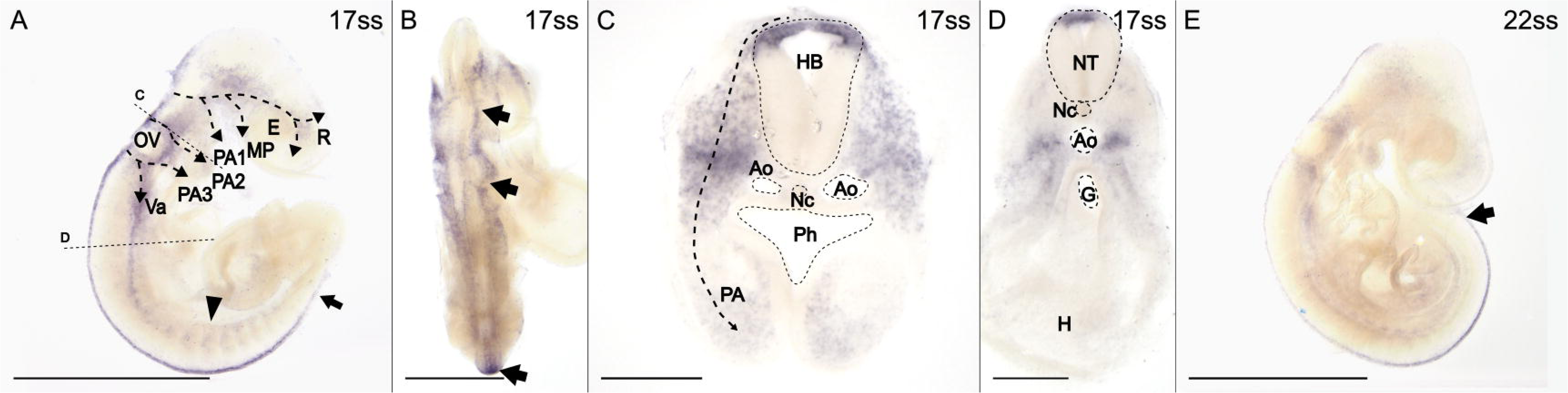
*Snai2* is expressed in both premigratory and migratory NCCs. Arrows indicates expression along the dorsal midline. **A-D** 17ss embryo in lateral (**A**) and dorsal (**B**) view and in cross-sections (**C** and **D**) at the levels indicated in **A**. Arrowhead indicates migration through the anterior half of the somites and dashed arrows indicate the inferred streams of migratory cranial NCCs. **E** 22ss embryo in lateral view. **Abbreviations:** Ao, aorta; E, eye; G, gut; H, heart; HB, hind brasin; MP, maxillary process; Nc, notochord; NT, neural tube; OV, otic vesicle; PA, pharyngeal arch; Ph, pharynx; R, rostrum; Va, vagal NCCs. **Scale** bar in **B** is 0.5 mm and scale bars in **C** and **D** are 0.1 mm. All other scale bars are 1 mm.

Similar to HNK-1, *Snai2* expression in putative premigratory NCCs along the dorsal midline of the trunk progressively extends caudally as development progresses (arrows in Figure 3A and E). *Snai2* expression is also evident in multiple streams of migratory NCCs traversing the somites (Figure A).

Taken together, the expression patterns of *Wnt1* and *Snai2* are compatible with the expected location of premigratory NCCs. As in chicken, but not mouse, *Snai2* seems to be expressed where NCCs are expected to undergo EMT.^66^ Similar to HNK-1, *Snai2*, but not *Wnt1*, is expressed in migratory NCCs. In addition, *Snai2* expression labels putative vagal NCCs more clearly than HNK-1.

### *Twist1*, *Snai1* and *Sox9* are expressed in NCCs biased towards mesenchymal fate

*Twist1* is a well-known NCC marker and transcription factor that has been studied in various other organisms. It has been found to be involved in EMT,^67^ maintenance of a migratory state, repression of neuronal differentiation,^68^ and induction of chondrocyte differentiation.^48^

Expression of *Twist1* is observed in all investigated stages (7 stages ranging from 13 through 51ss; Figure 4A-D). Cranial *Twist1* expression is missing along the dorsal midline but is consistent with the expected location of migratory NCCs (including the vagal branch) observed with HNK-1 and *Snai2* (dashed arrows in Figure 4A and C). The lack of *Twist1* expression in the presumed premigratory NCCs in the dorsal midline is consistent with findings in mouse and chicken,^68,69^ but different from frog and zebrafish.^67^ Later expression is mostly restricted to the maxillary process and pharyngeal arches (arrow and arrowhead in Figure 4D). Given the described role of *Twist1* as an inducer of chondrocytes,^43^ this expression likely label the formation of craniofacial structures in these regions. At early stages, there is also broad but transient expression of *Twist1* in the head ectoderm. While *Twist1* is also extensively expressed in the trunk, it is restricted to putative mesodermal cells in the somites and limb buds and will therefore be disregarded here.

**Figure 4.**
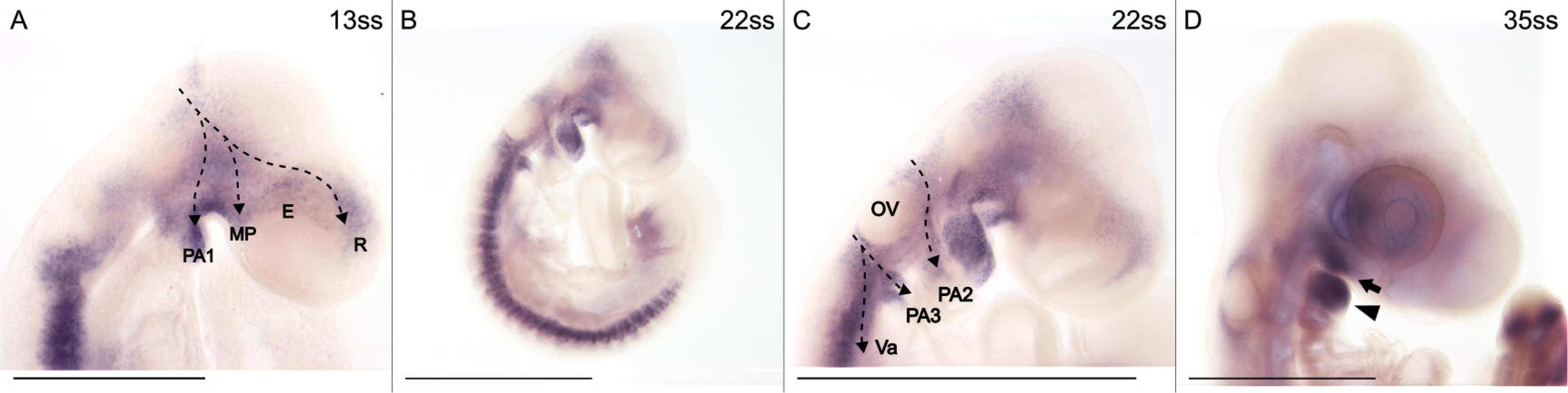
*Twist1* is expressed in migratory cranial NCCs. **A-C** Heads of 13, 17 and 22ss embryos in lateral view. Dashed arrows indicate the inferred streams of cranial NCC migration. **D** Head of 35ss embryo in lateral view. Arrow indicates the maxillary process and arrowhead indicates the first pharyngeal arch. **Abbreviations:** E, eye; MP, maxillary process; OV, otic vesicle; PA1-3, pharyngeal arch 1-3; R, rostrum; Va, posterior stream of vagal NCCs. **Scale bar** in **A** is 0.5 mm, all others are 1 mm.

*Snai1* is expressed in all investigated stages (7 stages ranging from 16 to 51ss) and its expression patterns resemble that of *Twist1* (Figure 5A-C). In contrast to *Snai2*, *Snai1* is not expressed where premigratory NCCs are expected to undergo EMT in the common wall lizard (Figure 5A and B), which is different from findings in mouse but consistent with findings in chicken.^66^ Compared to Twist1, the different streams of cranial migration are not as easily discernible for *Snai1* since the intensity of the expression signal is generally high (Figure 5A). Given this high intensity, it is possible that not only NCCs contribute to the signal.

**Figure 5.**
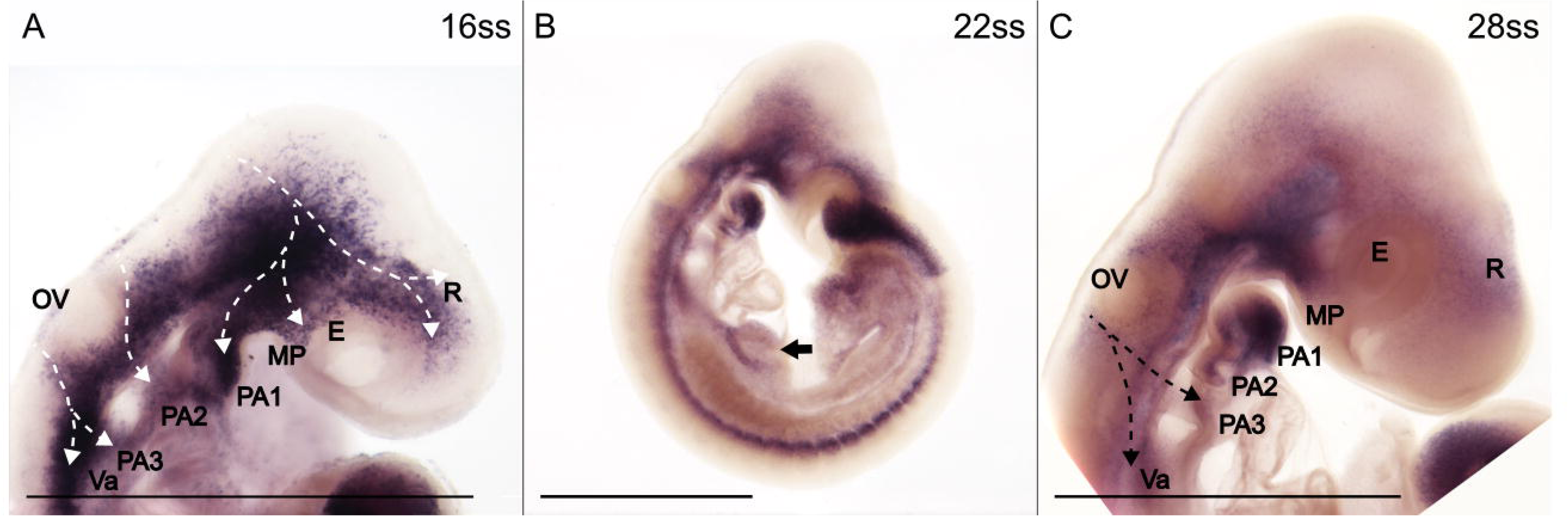
*Snai1* is expressed in migratory cranial NCCs. **A** Head of 16ss embryo in lateral view. Dashed lines indicate inferred streams of cranial NCC migration. **B** 22ss embryo in lateral view. Arrow indicates a bifurcation in the inferred caudal stream of vagal NCCs. Expression in the trunk is in tissues of non-NCC origin. **C** Head of 28ss embryo in lateral view. The dashed arrows indicate the third stream of inferred cranial NCC migration. **Abbreviations:** E, eye; MP, maxillary process; OV, otic vesicle; PA1-3, pharyngeal arch 1-3; R, rostrum; Va, posterior stream of vagal NCCs. **Scale bars** are 1 mm.

Together with *Snai1,* -*2* and *Twist1*, *Sox9* is known to be a crucial regulator of EMT^46,70^ and it is also known to regulate migration and chondrocyte differentiation of cranial NCCs.^46^ However, in the trunk, *Sox9* is expressed in premigratory but not migratory NCCs in chicken.^71^

In the common wall lizard, *Sox9* is expressed in all investigated stages (8 stages ranging from 13ss through 51ss; Figure 6A-D). In the head, *Sox9* appears to label both premigratory (arrows in Figure 6A) and migratory (dotted arrows in Figure 6A) NCCs. In the third stream, it is mainly the caudally directed putative vagal branch that can be inferred. Later in development, the expression in the inferred streams extends to much of the surface of the head (Figure 6C). At this point, the expression pattern resembles that of late *Twist1* expression.

**Figure 6.**
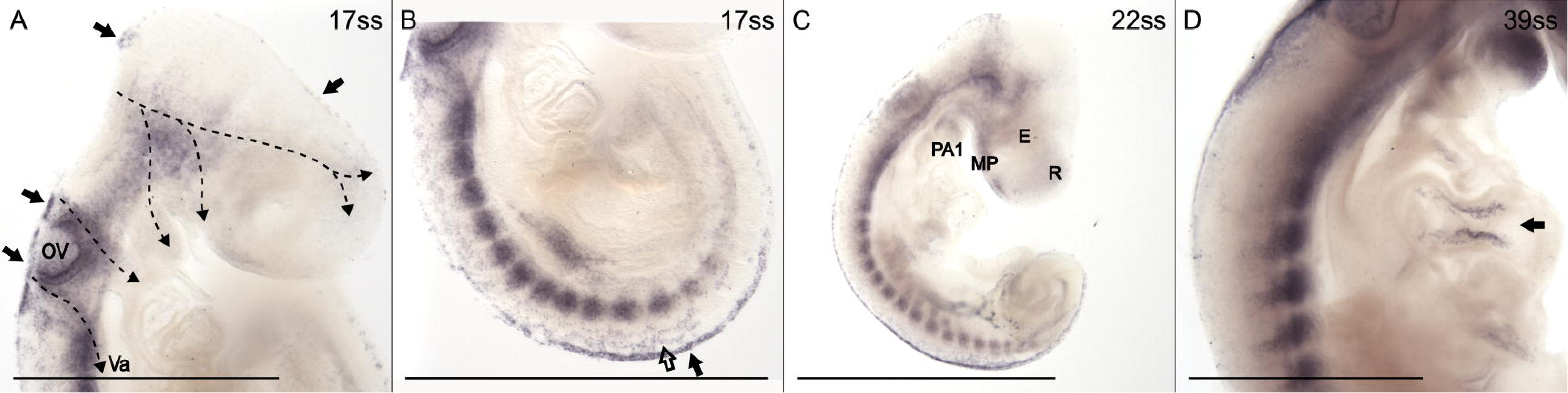
*Sox9* expression labels both premigratory NCCs and migratory cranial NCCs. **A, B** Head (**A**) and trunk (**B**) of 17ss embryo in lateral view. Dashed arrows indicate inferred streams of cranial NCC migration. Arrows indicate expression in the dorsal midline and open arrow indicates early migratory NCCs. **C** 22ss embryo in lateral view. **D** Trunk of 39ss embryo in lateral view. Arrow indicates expression in the heart. **Abbreviations:** E, eye; MP, maxillary process; OV, otic vesicle; PA, pharyngeal arch; R, rostrum; Va, posterior stream of vagal NCCs. Scale bars in **A** and **B** are 0.5 mm, all others are 1 mm.

*Sox9* expression in the trunk is consistent with premigratory NCCs along the dorsal midline, resembling that of *Snai2*. There is also expression in what appears to be early migratory NCCs just lateral to the dorsal midline that have not yet condensed into migratory streams (Figure 6B and C). Otherwise, *Sox9* does not appear to be expressed in migratory NCCs in the trunk, which is consistent with findings in chicken.^71^ There is expression in non-NCC related structures (Figures 6B and C), which will not be described in more detail here. Some *Sox9* expression can be seen in the heart at 39ss (Figure 6D) where it stains a structure that may be the outflow tract, which is partially derived from NCCs in mouse.^68^

In summary, the expression patterns of *Snai1*, *Twist1* and *Sox9* generally support the spatiotemporal location of migratory cranial neural crest cells inferred from HNK-1. In addition, they distinctly label regions such as the maxillary process and pharyngeal arches where facial skeleton will develop. *Sox9* is also expressed in the heart and in putative premigratory NCCs along the dorsal midline of both head and trunk.

### *Sox10*, *FoxD3* and *Phox2b* are expressed in NCCs biased towards neuronal and glial fates

After HNK-1, *Sox10* is arguably the most canonical NCC marker, and together with *FoxD3* and *Ets1* it is considered a NCC specifier gene.^46^ It is a marker of migratory NCCs,^72^ and it is necessary for NCC differentiation into melanocytes, neurons and glia.^46^

*Sox10* is expressed in all investigated stages (11 stages ranging from 13 to 64ss; Figure 7 A-P). All three streams of migratory cranial NCCs inferred from the HNK-1 staining are clearly expressing *Sox10* (Figure 7A and B). However, *Sox10* expression, in contrast to HNK-1, *Snai2, Snai1, Twist1* and *Sox9* does not stain the streams all the way to the rostrum, maxillary process and pharyngeal arches (Figure7A). As development progresses, cranial *Sox10* expression, more distinctly than HNK-1 staining, localizes in some cranial placodes (arrow in Figure 7H-J) and subsequently the corresponding nerves of the cranial peripheral nervous system (Figure 7L, M and P). A putative vagal stream migrating caudally in the trunk from the third cranial stream is marked by *Sox10* expression (arrowhead in Figure 7H and J). Later in development, the vagus nerve is clearly discernable through *Sox10* expression (Figure 7P).

**Figure 7.**
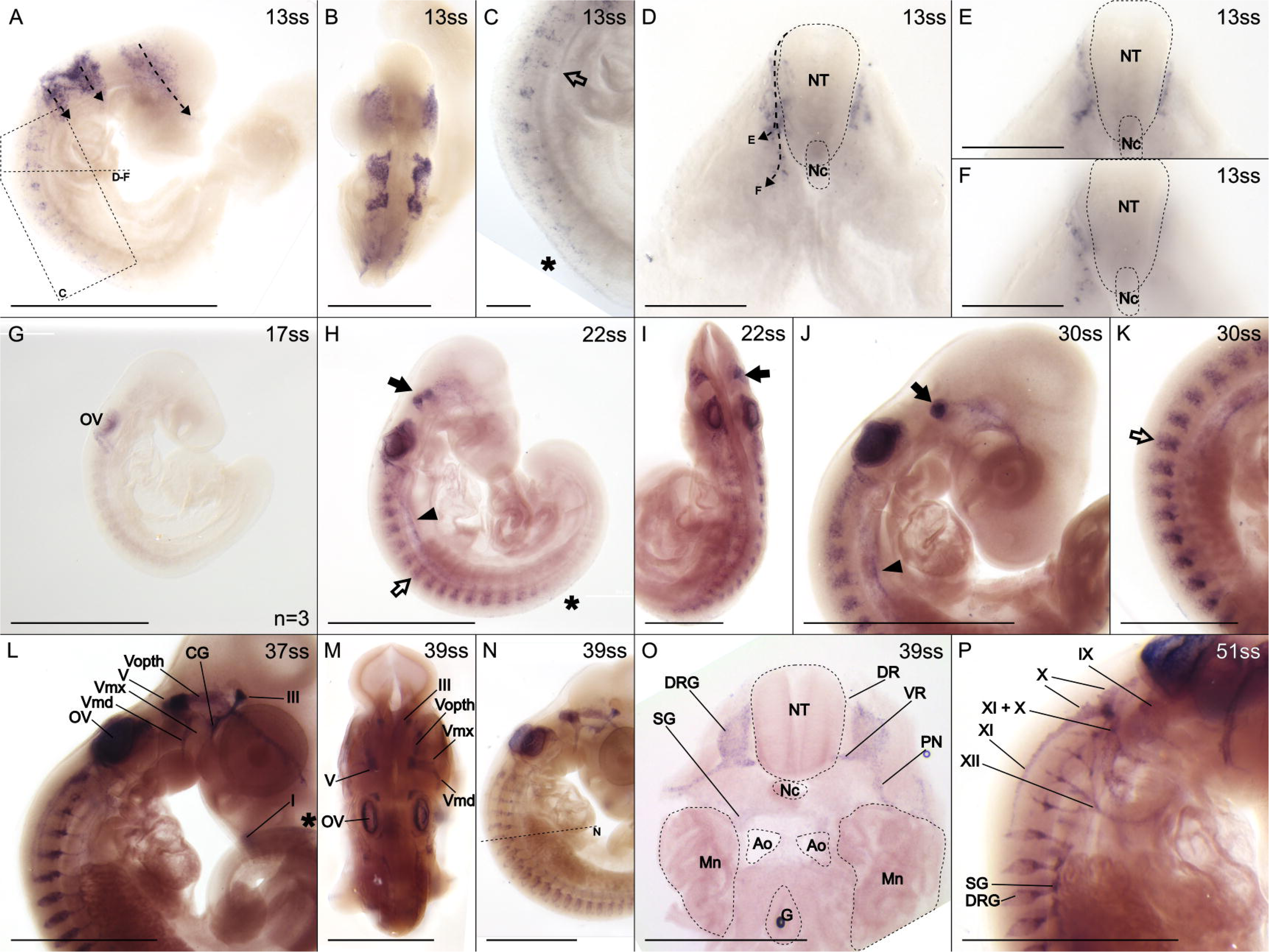
*Sox10* expression identifies migratory NCCs and the developing peripheral nervous system. Asterisks indicate premigratory and early migratory NCCs along the dorsal midline, arrows indicate the trigeminal nerve placode, open arrows indicate migratory trunk NCCs and arrowheads indicate caudally migrating vagal NCCs. **A-F** 13ss embryo in lateral (**A, C**) and dorsal (**B**) view and in cross-section (**D**-**F**). The positions of **C-F** are indicated in **A**. Dashed arrows indicate inferred streams of NCC migration. **D** is a composite of many focal planes, while **E** and **F** are single focal planes. **G** 17ss embryo in lateral view. Notice the weak signal, a result replicated in three separate experiments. **H-I** 22ss embryo in lateral (**H**) and dorsal (**I**) view. **J**-**K** Head (**J**) and trunk (**K**) of 30ss embryo in lateral view. **L** 37ss embryo in lateral view. Asterisk indicates weak expression along the dorsal midline of the tail. **M**-**O** 39ss embryo in dorsal (**M**) and lateral (**N**) view, and in cross-section (**O**). The approximate level of the cross-section in **O** is indicated by a dashed line in **N. P** 51ss embryo in lateral view. **Abbreviations:** Ao, aorta; CG, ciliary ganglion; DR, dorsal root; DRG, dorsal root ganglion; G, gut; Mn, mesonephros; Nc, notochord; NT, neural tube; OV, otic vesicle; PN, peripheral nerve; SG, sympathetic ganglion; VR, ventral root. **Cranial nerves:** I, olfactory nerve (also known as cranial nerve 1); III, oculomotor nerve (also known as cranial nerve 3); V, trigeminal nerve (also known as cranial nerve 5); Vopth, ophthalmic branch of the trigeminal nerve; Vmd, mandibular branch of the trigeminal nerve; Vmx, maxillary branch of the trigeminal nerve; IX, glossopharyngeal nerve (also known as cranial nerve 9); X, vagus nerve (also known as cranial nerve 10); XI, spinal accessory nerve (also known as cranial nerve 11); XII, hypoglossal nerve (also known as cranial nerve 12). **Scale bars** in **B** and **O** are 0.5 mm and **C**, **D**, **E** and **F** are 0.1 mm. All other scale bars are 1 mm.

Early *Sox10* expression in the trunk is largely similar to the spatiotemporal distribution of NCCs inferred from HNK-1 staining including putative premigratory and migratory NCCs (Figure 7A, C, H and K). However, the expression in the putative premigratory NCCs along the dorsal midline is rather weak (asterisks in Figure 7C, H and K). In cross-section at the level of the heart at 13ss, the inferred streams are revealed to migrate ventrally along the edge of the neural tube (Figure 7D-F). By looking at different levels along the anterior-posterior axis, two different routes of migration can be distinguished; one turns laterally and enters the somite (Figure 7D and E), presumably give rise to dorsal root ganglia.^18^ The other one continues ventrally (Figure 7D and F), likely giving rise to sympathetic ganglia.^18^ The expression in the migratory streams subsequently condenses to a row of dorsal root ganglia in each somite with ventral projections to the much smaller ganglia of the sympathetic chain (Figure 7L). The dorsal root ganglia, including their dorsal and ventral roots and peripheral projections, are clearly visible in cross-section, while sympathetic ganglia are evident more ventrally, though less strongly expressing *Sox10* (Figure 7O).

*FoxD3* is an important transcription factor in NCC development.^17^ Early in NCC development, *FoxD3* acts as an activator, allowing NCCs to delaminate from the neural tube, but later its role changes and it maintains the multipotency of NCCs by repressing differentiation.^18,73,74^

In the common wall lizard, *FoxD3* is expressed in all investigated stages (9 stages from 13 to 64ss), and its expression patterns are similar to those of *Sox10*, although mostly not as strong. Like *Sox10, FoxD3* stain premigratory NCCs along the dorsal midline (Figure 8A-C), migratory NCCs including three cranial streams (Figure 8A and B), and NCCs condensing in the peripheral nervous system of both the head and trunk (Figure 8 C and D). Unlike *Sox10*, *FoxD3* staining in premigratory NCCs along the dorsal midline is strong. It also stains in the geniculate ganglion (open arrow in Figure 8D) but not the otic vesicle.

**Figure 8.**
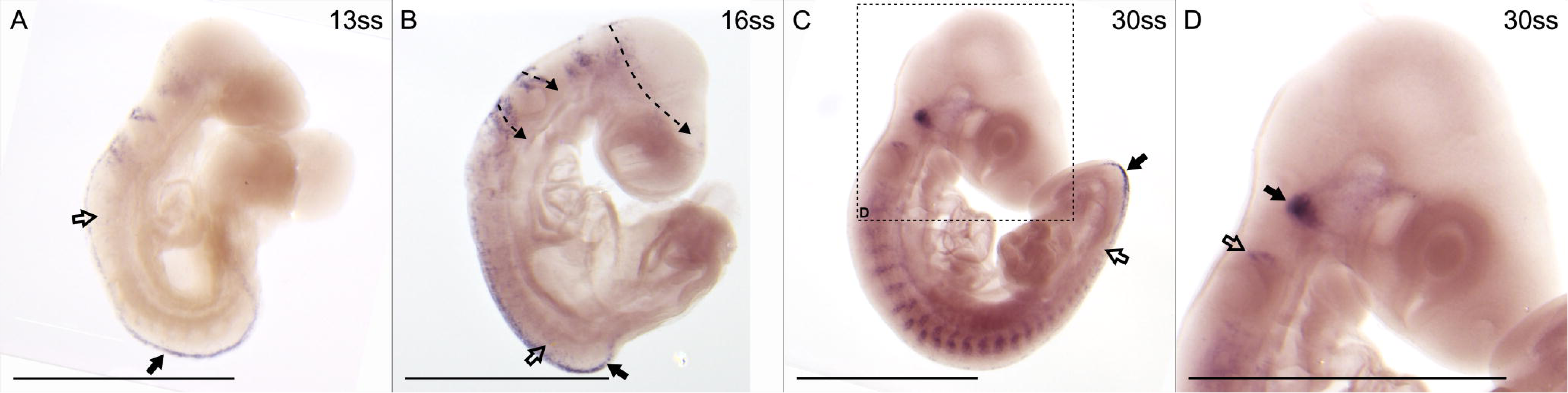
*FoxD3* expression labels both premigratory and migratory NCCs as well as the developing peripheral nervous system. **A** and **B** 13 and 16ss embryos in lateral view. Dashed arrows indicate streams of migratory cranial NCCs, arrows indicate expression in premigratory NCCs and open arrows indicate early migratory NCCs. **C** and **D** 30ss embryo in lateral view. Arrow indicates the trigeminal nerve placode and open arrow indicates the geniculate ganglion (also known as cranial nerve ganglion 7) The position of **D** is indicated in **A**. **Scale bars** are 1 mm.

Downstream of *Sox10, Phox2b* is a regulator of NCC differentiation into autonomic and especially sympathetic neurons and chromaffin cell progenitors.^46,75–77^ In the common wall lizard, *Phox2b* is expressed in all investigated stages (7 stages ranging from 17-51ss) except the youngest (17ss; data not shown). However, the expression is most distinct at 39ss (Figure 9A-C), and expression before and after this stage is generally restricted and weak. *Phox2b* expression localizes to several of the cranial ganglia, which are co-derived from the cranial placodes and NCCs.^78^ There is also significant expression in the hindbrain in the form of three longitudinal stripes on each side (Figure 9A), similar to previous results in mouse.^79^ Contrary to previous results in mouse and chicken there is no distinct expression of *Phox2b* in the dorsal root ganglia in any of the investigated stages.^78,79^

**Figure 9.**
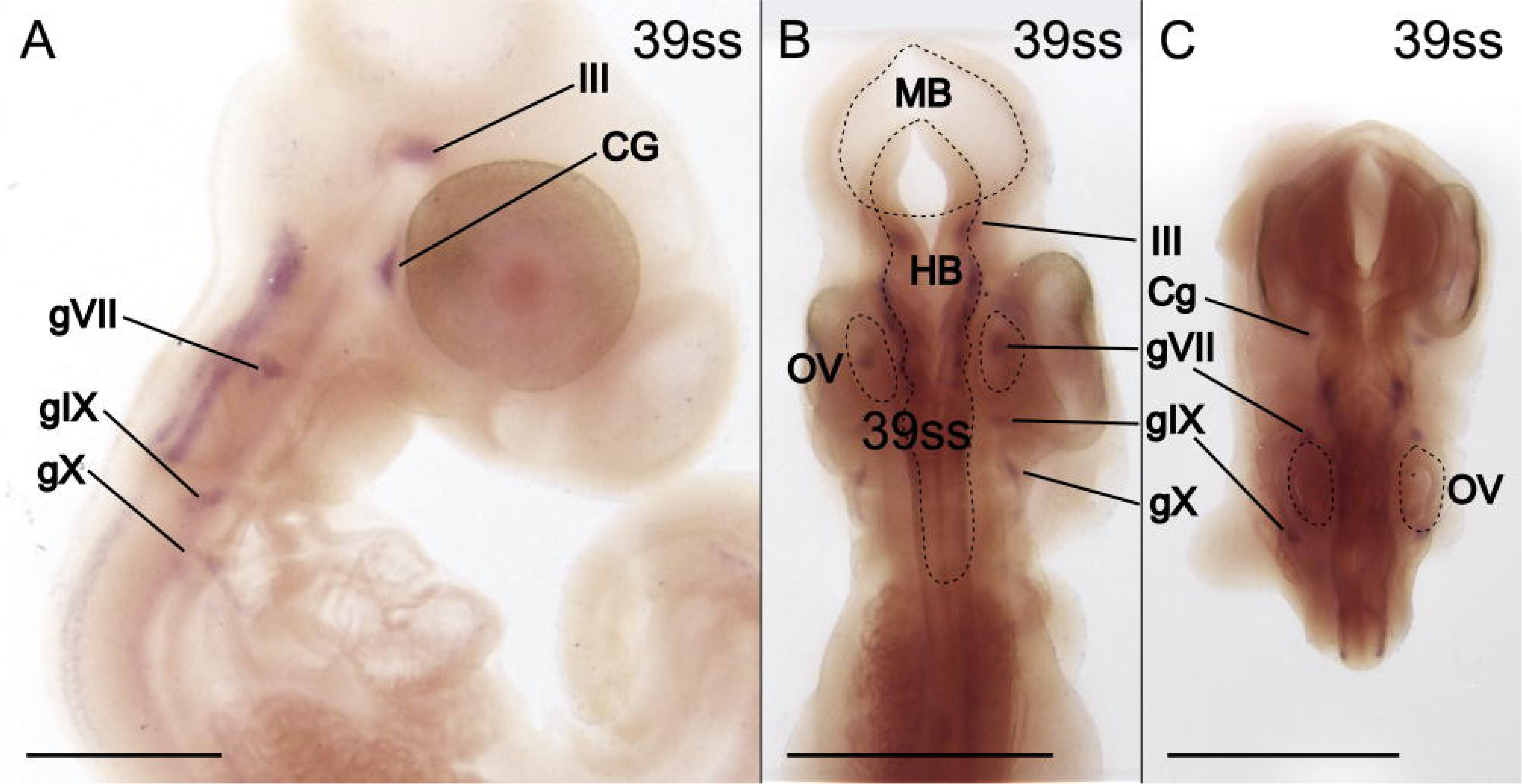
*Phox2b* expression identifies cranial ganglia. **A-C** Head of 39ss embryo in lateral (**A**) and dorsal (**B** and **C**) view. Expression signals in several cranial nerve ganglia are indicated. **Abbreviations:** CG, ciliary ganglion; III, oculomotor nerve (also known as cranial nerve 3); gVII, geniculate ganglion (also known as cranial nerve ganglion 7); IX, glossopharyngeal ganglion (also known as cranial nerve 9); X, vagus nerve (also known as cranial nerve 10); OV, otic vesicle. **Scale bars** are 1 mm.

Taken together, gene expression profiles of NCCs biased towards neuronal and glial fates (*Sox10*, *FoxD3* and *Phox2b*) largely confirm and expand the spatiotemporal distribution of NCCs inferred from HNK-1 staining. Together their expression patterns clearly visualize the contribution of NCCs to the development of the peripheral nervous system, including the sympathetic nervous system.

### *Th* expression identifies NCCs differentiating into chromaffin cells

In mouse, *Tyrosine hydroxylase* (*Th*) has been identified as a suitable marker for separating the chromaffin cells from the sympathetic neurons.^76,80^ It is an enzyme catalyzing one of the reactions in the metabolic pathway that result in the production of adrenalins.^81^

*Th* expression is first detected at 35ss and is maintained at least until 51ss. The expression is located internally in the trunk, at the level of the forelimb bud (Figure 10A-F). *Th* expression is localized in two lateral parallel lines (Figure 10B) in the dorsomedial corner of the mesonephros (Figure 10D). This is putatively the position where the sympathetic ganglia including the adrenal glands are developing. The areas of expression are discontinuous (see for example arrowheads in Figure 10E) concordant with chromaffin cells being restricted to the sympathetic ganglia including the adrenal glands.

**Figure 10.**
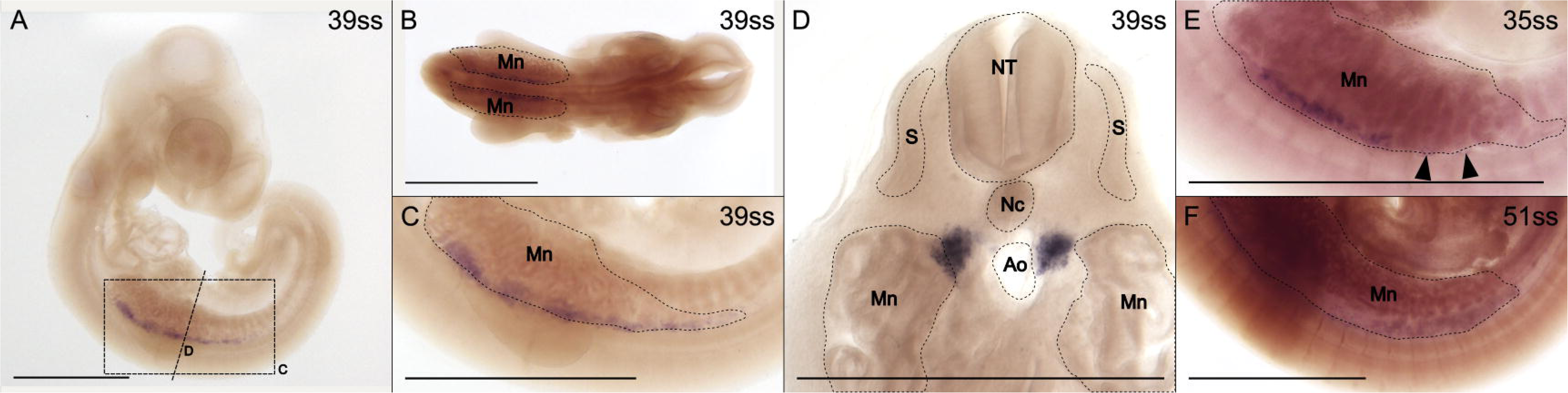
*Th* expression identifies the chromaffin cells of the putative adrenal gland. **A-D** 39ss embryo in lateral (**A, C**) and dorsal (**B**) view and in cross-section (**D**). Dashed rectangle and line in **A** indicate the approximate position of panels **C** and **D** respectively. **E, F** Trunk of 35 and 51ss embryos in lateral view. Arrowheads indicate discontinuity in the expression. **Abbreviations:** Ao, aorta; Mn, mesonephros; Nc, notochord; NT, neural tube; S, somite. **Scale bar** in **D** is 0.5 mm, all others are 1 mm.

## Conclusions

Taken together, the insights gained from HNK-1 immunostainings and gene expression patterns of nine marker genes provide a detailed profile of the spatiotemporal distribution of NCCs in developing wall lizard embryos (Figure 11). By the time of egg-laying, the embryo has already reached at least 13ss at which extensive migration is ongoing in the head. Around 35ss, most migration is completed and differentiation towards various fates is well underway. In the following, we briefly summarize the major patterns of NCC development at four different stages. At 14ss, premigratory NCCs are present along the dorsal neural tube all the way from the midbrain-hindbrain boundary to the last formed somite pair (orange in Figure 11A). Streams of cranial NCCs are already migratory. In the caudal trunk, there is no migration observed at this stage. At more anterior regions in the trunk, a field of NCCs lateral to the dorsal midline indicates that the first wave of NCC migration has started but these cells have not yet entered the somites (green in Figure 11A). In the anterior-most part of the trunk, migration has proceeded further with the first migratory streams traversing the anterior parts of the somites (turquoise in Figure 11A).

**Figure 11.**
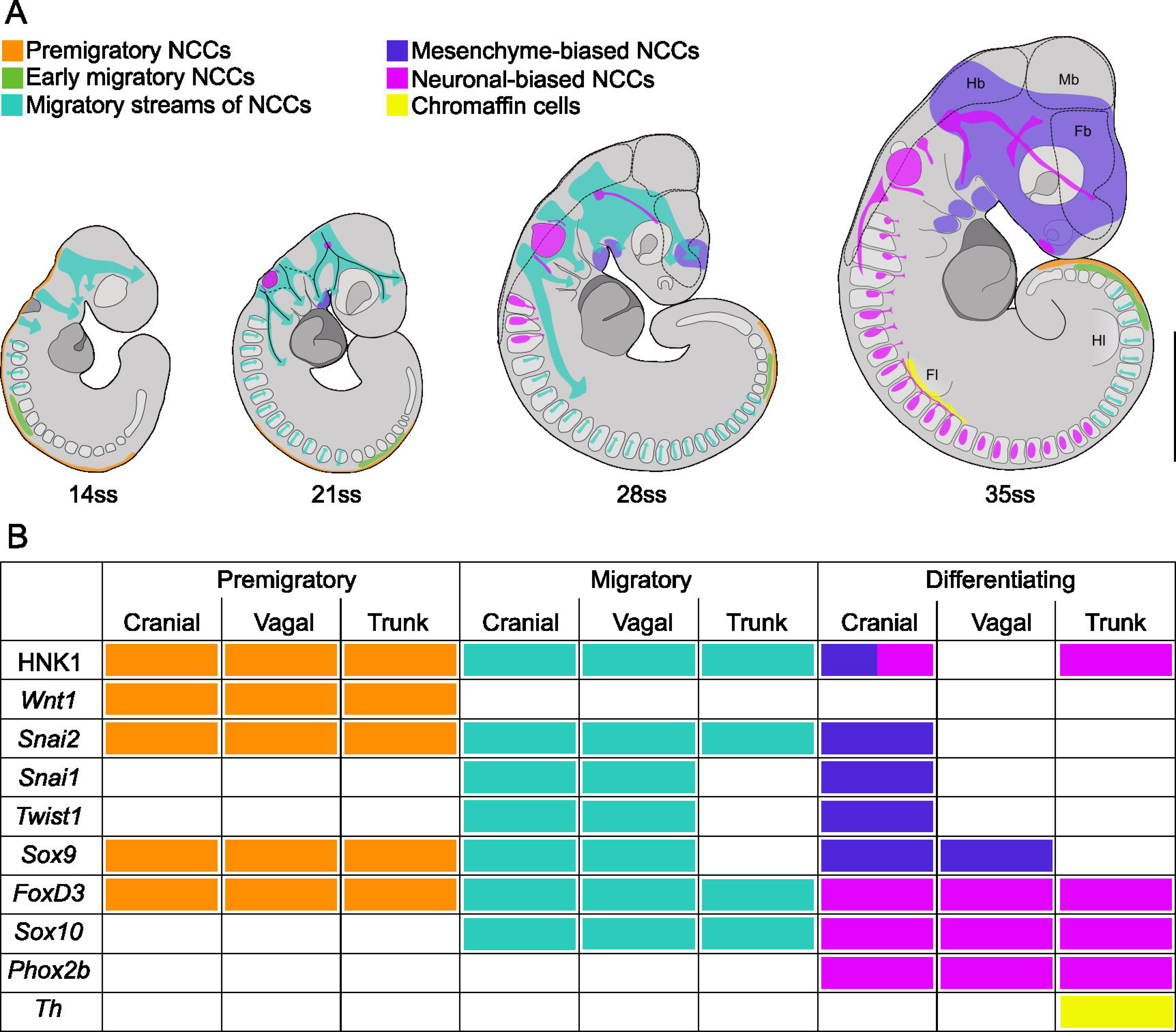
Synthesis of the spatiotemporal distribution of NCCs in developing common wall lizards. **A** Summary of the inferred distribution and migratory streams of NCCs in four developmental stages of the common wall lizards. Different groups of NCCs are color-coded according to their state or expected fate. **B** Schematic overview of the state of NCCs (premigratory, migratory or differentiating) and their respective domain (cranial, vagal or trunk) with respect to their expression of individual marker genes and the presence of HNK-1 positive staining. **Abbreviations**: Fb, forebrain; Fl, forelimb; Hb, hindbrain; Hl, hindlimb; Mb, midbrain.

At 21ss, NCC specification and migration in the trunk has proceeded further posteriorly and migration now occurs through several of the anterior-most somites (approximately the first 13-15). In the head, the streams of migratory cranial NCCs have reached further and are bifurcating more. NCCs with mesenchymal fate are located in the first pharyngeal arch (purple in Figure 11A). and NCCs with neural fate are located in the otic vesicle and trigeminal nerve placode (magenta in Figure 11A). Vagal NCCs appear as a branch extending caudally through the anterior trunk from the third cranial stream.

By 28ss, NCC specification and migration in the trunk has proceeded even further posteriorly and the first differentiated dorsal root ganglia occur in the anterior-most somites (approximately the first 3; magenta in Figure 11A). The cranial streams are now more advanced. NCCs with mesenchymal fate appear in the maxillary process and the rostrum and NCCs with neural fate are expanding in cranial placodes and the first neural projections (magenta in Figure 11A).

At 35ss, specification and migration of NCCs is restricted to the caudal-most trunk region. Neuronal differentiation is well underway in both the dorsal root ganglia and the sympathetic ganglia of the trunk. At the level of the forelimb, some NCCs have differentiated into chromaffin cells (yellow in Figure 11A). The mesenchymal NCCs are now distributed broadly in the head including the first three pharyngeal arches, maxillary process, rostrum and close to the olfactory pit. Cranial nerves have developed further, the olfactory (I), oculomotor (III), trigeminal (V), glossopharyngeal (IX), vagus (X) and spinal accessory nerves are now all clearly distinguishable. This synthesis is derived from the collective information gathered from ten markers (Figure 11B), which proved to be crucial to overcome incomplete information stemming from individual markers. For example, HNK-1 immunostaining and *Phox2b* expression signals are not located in the dorsal root ganglia, yet other markers, for example *Sox10* and *FoxD3*, clearly identify this tissue as NCC-derived. The HNK-1 signal in dorsal root ganglia reported for the veiled chameleon^9^ is also less distinct than typically seen in other species. Thus, the lack or reduction of HNK-1 on dorsal root ganglia may be a more widespread feature of lizards. In addition, since there are no distinct NCC markers that identify exclusively NCCs, off-target staining necessarily needs to be taken into account. Examples include the expression of *Snai1* and *Twist1* in trunk mesoderm or the presence of HNK-1 on blood cells.

Comparing our synthesis of NCCs in the common wall lizard to those reported for other squamates (the Californian kingsnake^7^, Egyptian cobra^82^ and veiled chameleon^9^) reveals that NCC biology is largely conserved. One caveat for this comparison is that our analysis is restricted to embryos older than 13ss, and we therefore lack observations of the earliest stages of NCC specification and migration, particularly in the head. It was therefore not possible to investigate if common wall lizards, similar to chameleons^9^ and crocodiles^76^, possess an early rostral stream of migrating cranial NCCs stemming from the level of the forebrain. In future studies, efforts could be made to obtain earlier embryos by, for example, treating gravid females hormonally to induce premature egg laying,^84^ or by selecting lizard species that lay eggs at earlier developmental stages.

The two main exceptions of NCC regulation in squamates relative to other tetrapods that have been reported to date are the fourth stream of migrating NCCs from the hindbrain observed in the veiled chameleon^9^ (also found in an alligator and an ostrich)^16^ and the late wave of trunk NCCs taking a ventromedial (instead of the expected dorsolateral route), which was found in the Californian kingsnake.^7^ Our observations in the common wall lizard do not clearly identify a fourth stream of NCCs from the hindbrain. However, to confidently rule out the existence of this stream, further analyses including cell lineage tracing may be required. Regarding the ventromedial route of late wave trunk NCCs, we do find ventromedial intra-somitic migration that is coinciding with dorsolateral migration, but it is not clear that these correspond to the late wave ventromedial migration reported for the California kingsnake.^7^ On a more technical note, it would also be desirable to confirm and cross-validate results in various species with the same antibody given that observed species-level differences might be partially attributed to the use of different antibodies targeting HNK-1.

In summary, our study provides the first complete description of NCCs, including cranial, vagal and trunk NCCs, in a lacertid and contributes to our understanding of taxon-specific differences in the dynamic distribution of this cell type. Thus, it lays the foundation for future work targeting the role of this cell type in evolutionary diversifications.

## Experimental procedures

### Embryo collection

All embryos stem from captive breeding colonies of common wall lizards that were wild caught in central Italy. Parents showed variation in phenotypes (coloration, morphology, and behavior), including in traits derived from the neural crest. In the present study, we describe broad developmental patterns that we do not expect to differ on a population level, and we therefore do not discriminate between embryos of different parental origins.

Adult wall lizards were kept as described in Feiner et al., (2018).^85^ Eggs were collected during the breeding seasons (April-July) of 2021 to 2023. Each cage had a pot filled with moist sand in which females laid their clutches. The eggs were collected each morning, and one egg per clutch was dissected immediately to determine the developmental stage (somite count) of the embryos in the clutch. The rest of the eggs were incubated at 24°C in moist vermiculite until the appropriate stage was reached. We aimed to sample a distribution of stages from as young as possible up to around 60ss. Based on 563 eggs, the mean embryonic stage on the day of oviposition was estimated to be 21.4ss (sd = 5.2 somites) and the earliest embryos were at 11ss. The pace of somite formation is constant and has previously been estimated to be four somites per day when incubated at 24°C.^85^

Egg dissections were performed in cold, nuclease-free PBS (10mmol/L phosphate buffer, 2.7mmol/L KCl and 137mmol/L NaCl, pH 7.4) after which each embryo was immediately transferred to 4% paraformaldehyde in PBS for overnight fixation at 4°C on a rotating platform. After fixation, embryos were transferred to gradually increasing concentrations of methanol and stored in 100% methanol at −20°C. A small subset of embryos intended to be used in probe synthesis (see below) was dissected as described above, incubated in RNAlater (Invitrogen AM7020) at 4°C for 24h and then transferred to −80°C for storage.

### Immunohistochemistry

Paraformaldehyde-fixed embryos were rehydrated in PBS-T with decreasing concentrations of methanol. Embryos were blocked using TN-blocking buffer with bovine serum albumin (1% bovine serum albumin, 0.1mol/L Tris pH7.5, 0.15 mol/L NaCl in H_2_O) followed by overnight incubation with primary antibodies against HNK-1 (1:500; Anti-Hu CD57 eBioscience 11-057742) in TN-blocking buffer. Embryos were washed with PBS and incubated overnight with secondary antibodies (1:200; Alexa Flour 488 goat anti-mouse IgM (µ chain) Invitrogen A21042) in TN-blocking buffer. Embryos were washed in PBS and stained with DAPI (1:500; ThermoScientific 62248). Lastly, embryos were cleared by overnight incubation in RapiClear 1.49 (SUNJin Lab) and then mounted in fresh RapiClear medium between two 0.17µm cover slips using iSpacers (SUNJin Lab). The stained and cleared embryos were imaged using a confocal microscope (Leica TCS SP8) using a 20X/0.75 IMM objective (plan apochromatic multi-immersion). Tiles were merged and 2D images were rendered using the LAS X software version 0.5.7.23225.

### Probe synthesis for *in situ* hybridization

Total RNA was extracted from the RNAlater-stored embryos using the RNeasy Mini Kit (Qiagen) including DNase digestion and stored in nuclease-free water at −80°C. Extracted RNA was used as template in reverse transcription into cDNA using SuperScript III Reverse Transcriptase and oligo-dT primers following the manufacturer’s instructions. The selected marker genes were amplified by polymerase chain reaction (PCR) from the cDNA using GoTaq G2 Hot Start Green Master Mix (Promega), and primers designed using the PodMur_1.0 reference genome.^86^ Primers were designed to amplify 800-1200 bp of the gene of interest with approximately 500 bps in the 3’UTR and T7 and SP6 promoters attached to the reverse and forward primers, respectively (see Table 1 for primer sequences). Amplified gene fragments were Sanger-sequenced to confirm each amplicon’s identity. PCR products were purified using the MinElute PCR Purification Kit (Qiagen) and *in vitro* transcribed to produce antisense RNA-probes using the DIG Probe Synthesis Kit (Roche). The resulting RNA-probes were purified using the RNeasy Mini Kit (Qiagen) and stored in nuclease-free water at −80°C.

**Table 1.**
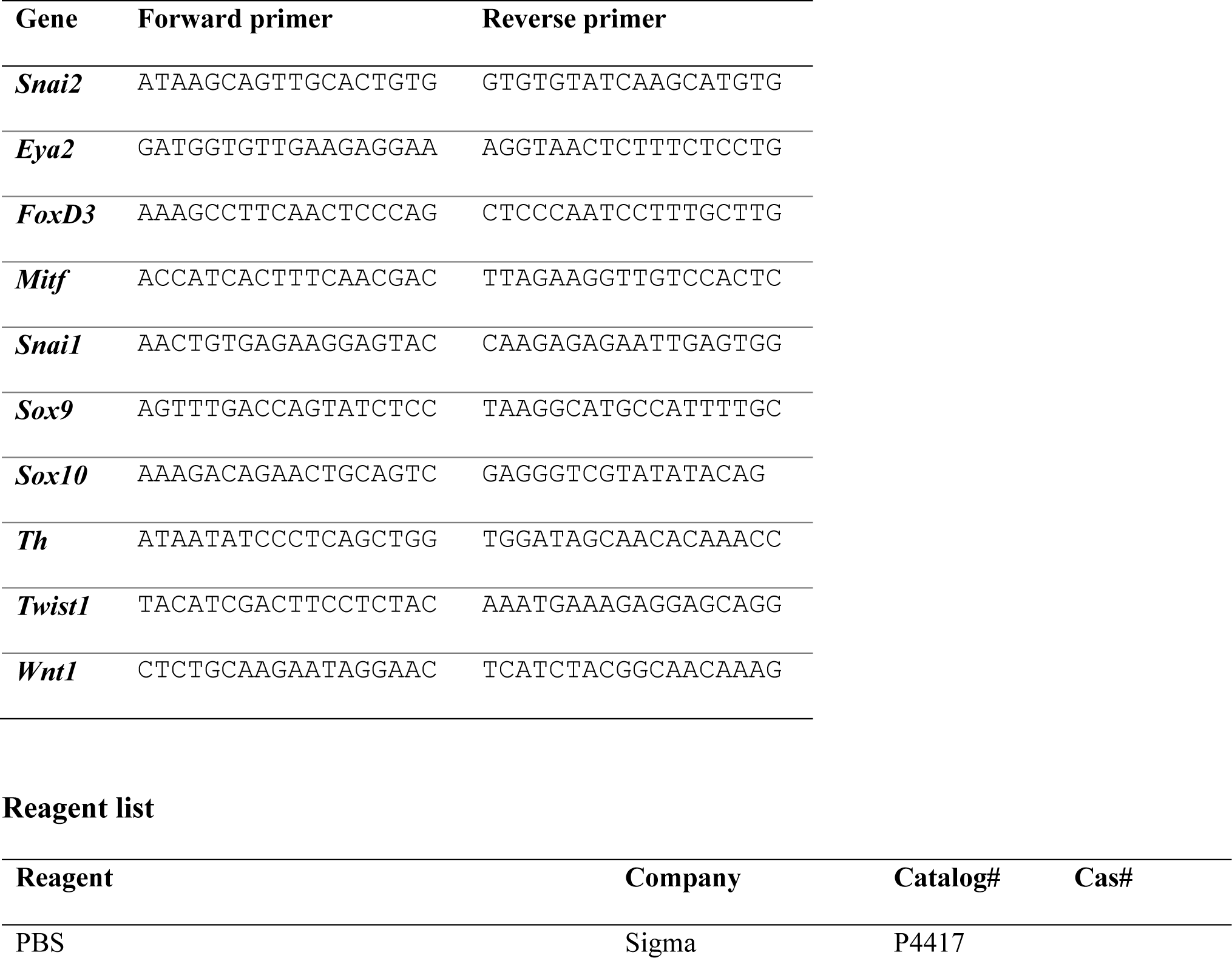
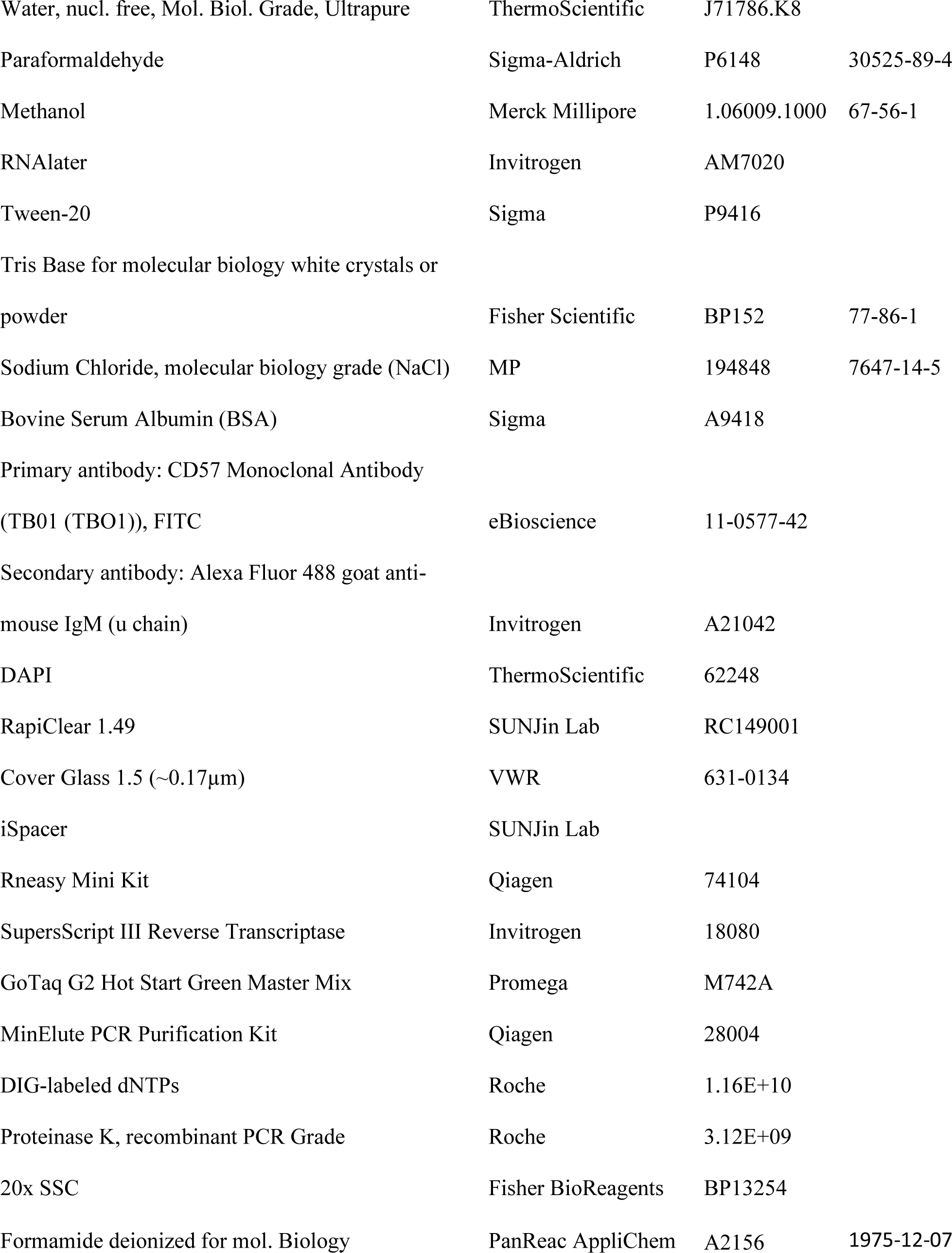
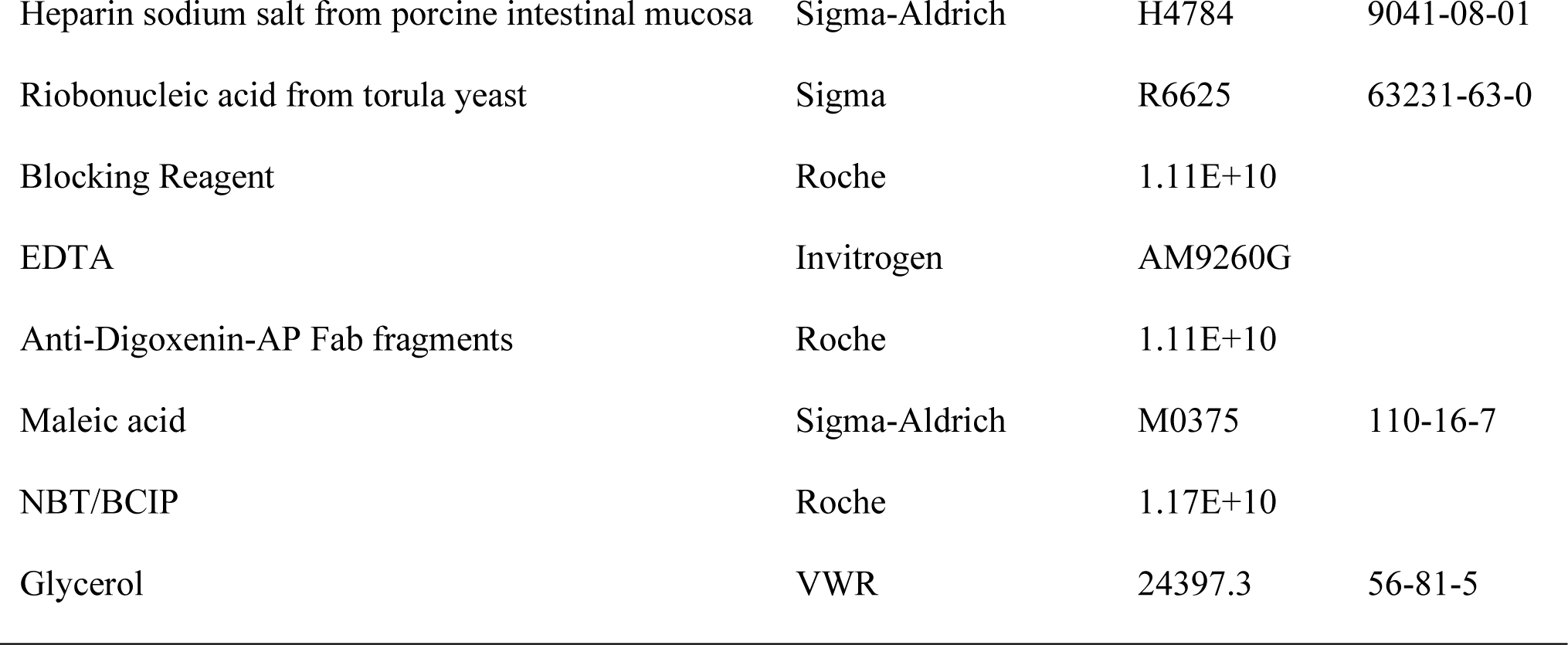
Sequences of primers used in this study (5’ to 3’).

### Whole mount *in situ* hybridization

Following the methodology in Feiner (2019)^87^ paraformaldehyde-fixed embryos were rehydrated in nuclease-free PBS-T (1x PBS with 0.1% Tween-20) using gradually lower concentrations of methanol. Embryos were permeabilized using Proteinase K, followed by refixation using 4% paraformaldehyde in PBS. Hybridization was done with 100 ng/mL probe at 68°C in hybridization buffer (5x SSC pH 7.5, 50% deionized formamide, 50 µg/ml heparin, 0.3 mg/ml torula RNA, 2% Tween-20, 2% blocking reagent and 10 mmol/L EDTA). Hybridized embryos were blocked in blocking buffer (1.5% blocking reagent in PBS-T), followed by incubation with 0.1875 U/mL anti-DIG antibody in blocking buffer. This was followed by 1.5 days of washes in maleic acid buffer (15 mmol/L maleic acid pH7.5, 300 mmol/L NaCl and 0.1% Tween-20) before color development with 2% NBT/BCIP in alkaline phosphatase buffer (0.1 mol/L Tris Base pH 9.5, 0.1mol/L NaCl, 1 mmol/L MgCl_2_ and 0.1% Tween-20). Once a good signal-to-noise ratio was achieved, the reaction was stopped by extensive washes in PBS-T and re-fixing in 4% paraformaldehyde. Some stained embryos were cross-sectioned by hand using a scalpel. Stained embryos and sections were photographed in glycerol using a Nikon SMZ18 stereo microscope with a Nikon SHR Plan Apo 2X WD: 20 objective and Nikon DS-Ri1 camera.

### Gene orthology

Since the orthology relationships of *Snai1* and *Snai2* to other tetrapod genes was unclear^66^ a gene tree was constructed to confirm their identities. Both sequences from the common wall lizard were obtained from Ensemble and blasted (protein blast) against the Human, mouse, and chicken databases on NCBI, and the top hits were retrieved. Alignment (using the muscle algorithm), model selection (JTT+G) and maximum likelihood tree construction was performed in MEGA (Figure 12).^88^

**Figure 12.**
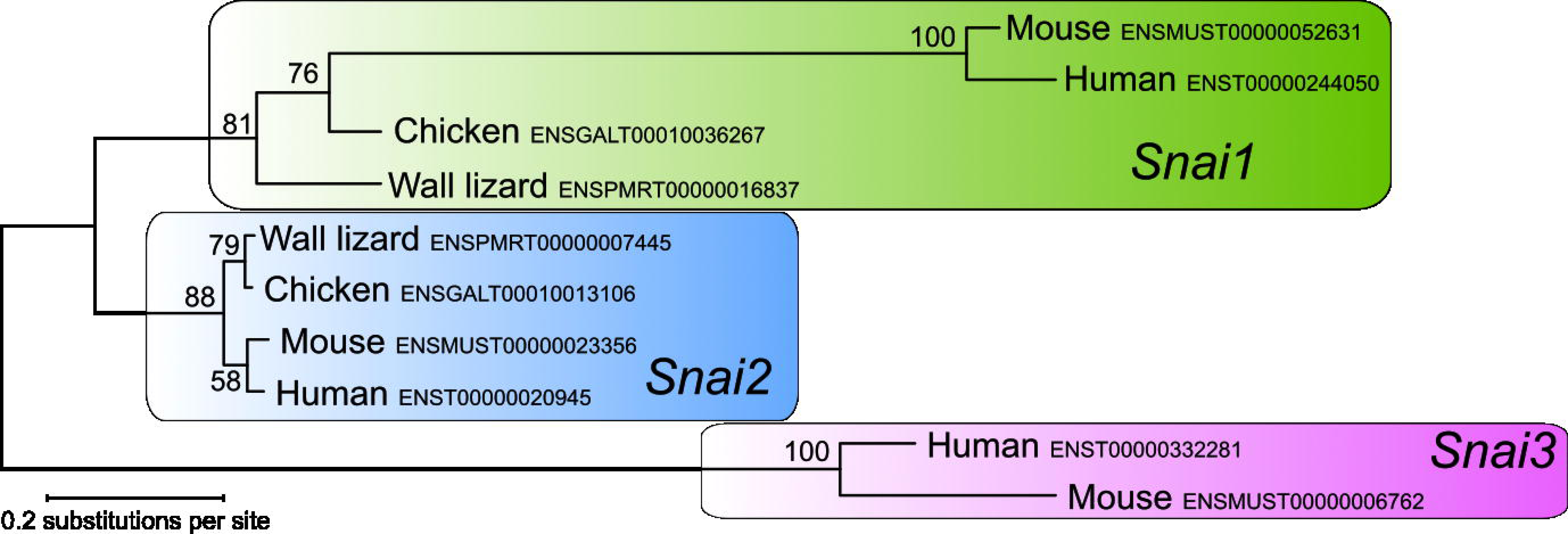
Gene tree of *Snai1*, *-2* and *-3*. Orthology relations of the transcription factors *Snai1* and *-2* in the common wall lizard was confirmed with regard to human, mouse and chicken. The *Snai3* gene appears to be absent from the common wall lizard genome. Accession numbers from Ensemble (wall lizard) and NCBI (mouse, human and chicken) are given next to each branch tip.

## Acknowledgements

We are grateful to Stanley Heinze, Michael Bok, Marjorie Lienard and Carina Rasmussen for technical advice and Ola Gustafsson for help with confocal microscopy. We thank Tobias Uller for valuable discussion and help with animal husbandry. We thank two anonymous reviewers for insightful comments.

## Data availability statement

Not applicable.

## Ethics approval statement

The embryos were collected from eggs laid by animals kept under breading permit 5.2.18-10993/18 granted by the Swedish ministry of agriculture.

## Patient consent statement

Not applicable

## Permission to reproduce material from other sources

Not applicable.

## References

1. Barresi MJF, Gilbert SF. Developmental Biology. 12th ed. Oxford, New York: Oxford University Press; 2020.

2. Alkobtawi M, Monsoro-Burq AH. The neural crest, a vertebrate invention. In: Evolving Neural Crest Cells. Evolutionary cell biology. Boca Raton: CRC Press; 2020:5–66. DOI: 10.1201/b22096

3. Abitua PB. The Hunt for Neural Crest in Invertebrate Chordates. In: Evolving Neural Crest Cells. Evolutionary cell biology. Boca Raton: CRC Press; 2020:137–156. 10.1201/b22096

4. Kuratani S. Neural Crest and Craniofacial Evolution of Early Vertebrates. In: Evolving Neural Crest Cells. Evolutionary cell biology. Boca Raton: CRC Press; 2020:185–218. 10.1201/b22096

5. Yu JK, Su YH. The Evolution of the Neural Border and Peripheral Nervous System—Insights from Invertebrate Deuterostome Animals. In: Evolving Nerual Crest Cells. Evolutionary cell biology. Boca Raton: CRC Press; 2020:103–136. 10.1201/b22096

6. Cerrizuela S, Vega-Lopez GA, Méndez-Maldonado K, Velasco I, Aybar MJ. The crucial role of model systems in understanding the complexity of cell signaling in human neurocristopathies. WIREs Mechanisms of Disease. 2022;14(1):e1537. 10.1002/wsbm.1537

7. Reyes M, Zandberg K, Desmawati I, De Bellard ME. Emergence and migration of trunk neural crest cells in a snake, the California Kingsnake (Lampropeltis getula californiae). BMC Developmental Biology. 2010;10. 10.1186/1471-213X-10-52

8. Khannoon ER, Alvarado C, Poveda R, de Bellard ME. Description of trunk neural crest migration and peripheral nervous system formation in the Egyptian cobra Naja haje haje. Differentiation. 2023;133:40–50. 10.1016/j.diff.2023.06.002

9. Diaz RE, Shylo NA, Roellig D, Bronner M, Trainor PA. Filling in the phylogenetic gaps: Induction, migration, and differentiation of neural crest cells in a squamate reptile, the veiled chameleon (Chamaeleo calyptratus). Developmental Dynamics. 2019;248(8):709–727. 10.1002/dvdy.38

10. Nichols DH. Neural crest formation in the head of the mouse embryo as observed using a new histological technique. Development. 1981;64(1):105–120. 10.1242/dev.64.1.105

11. Nichols DH. Formation and distribution of neural crest mesenchyme to the first pharyngeal arch region of the mouse embryo. American Journal of Anatomy. 1986;176(2):221–231. 10.1002/aja.1001760210

12. Serbedzija GN, Bronner-Fraser M, Fraser SE. Vital dye analysis of cranial neural crest cell migration in the mouse embryo. Development. 1992;116(2):297–307. 10.1242/dev.116.2.297

13. O’Rahilly R, Müller F. The development of the neural crest in the human. Journal of Anatomy. 2007;211(3):335–351. 10.1111/j.1469-7580.2007.00773.x

14. Le Douarin N, Kalcheim C. The Neural Crest. 2nd ed.; 1999. Accessed May 2, 2024. https://www.cambridge.org/se/academic/subjects/life-sciences/cell-biology-and-developmental-biology/neural-crest-2nd-edition, https://www.cambridge.org/se/academic/subjects/life-sciences/cell-biology-and-developmental-biology

15. Le Douarin NM. The avian embryo as a model to study the development of the neural crest: a long and still ongoing story. Mechanisms of Development. 2004;121(9):1089–1102. 10.1016/j.mod.2004.06.003

16. Kundrát M. Heterochronic shift between early organogenesis and migration of cephalic neural crest cells in two divergent evolutionary phenotypes of archosaurs: Crocodile and ostrich. Evolution and Development. 2009;11(5):535–546. 10.1111/j.1525-142X.2009.00352.x

17. Bronner-Fraser M. Analysis of the early stages of trunk neural crest migration in avian embryos using monoclonal antibody HNK-1. Developmental Biology. 1986;115(1):44–55. 10.1016/0012-1606(86)90226-5

18. Harris ML, Erickson CA. Lineage specification in neural crest cell pathfinding. Developmental Dynamics. 2007;236(1):1–19. 10.1002/dvdy.20919

19. Weston JA. 6 Sequential Segregation and Fate of Developmentally Restricted Intermediate Cell Populations in the Neural Crest Lineage. In: Bode HR, ed. Current Topics in Developmental Biology. Vol 25. Academic Press; 1991:133–153. 10.1016/S0070-2153(08)60414-7

20. Kuo BR, Erickson CA. Vagal neural crest cell migratory behavior: A transition between the cranial and trunk crest. Developmental Dynamics. 2011;240(9):2084–2100. 10.1002/dvdy.22715

21. Brandon AA, Almeida D, Powder KE. Neural crest cells as a source of microevolutionary variation. Seminars in Cell & Developmental Biology. Published online June 16, 2022. 10.1016/j.semcdb.2022.06.001

22. Darwin C. The Variation of Animals and Plants under Domestication. Vol 2. 2nd ed. London: John Murray; 1888.

23. Wilkins AS, Wrangham RW, Tecumseh Fitch W. The “domestication syndrome” in mammals: A unified explanation based on neural crest cell behavior and genetics. Genetics. 2014;197(3):795–808. 10.1534/genetics.114.165423

24. San-Jose LM, Roulin A. On the Potential Role of the Neural Crest Cells in Integrating Pigmentation Into Behavioral and Physiological Syndromes. Frontiers in Ecology and Evolution. 2020;8:278. 10.3389/fevo.2020.00278

25. Uller T, Moczek AP, Watson RA, Brakefield PM, Laland KN. Developmental Bias and Evolution: A Regulatory Network Perspective. Genetics. 2018;209(4):949–966. 10.1534/genetics.118.300995

26. Böhme W. Handbuch der Reptilien und Amphibien Europas Band2/II Echsen III (Podarcis). Wiesbaden: AULA-Verlag; 1986.

27. Eimer T, Eimer A. Untersuchungen Über Das Variiren Der Mauereidechse : Ein Beitrag Zur Theorie von Der Entwicklung Aus Constitutionellen Ursachen, Sowie Zum Darwinismus. Berlin: Nicolaische Verlags-Buchhandlung; 1881. 10.5962/bhl.title.49215

28. Kammerer P. Der Artenwandel Auf Inseln Und Seine Ursachen Ermittelt Durch Vergleich Und Versuch an Den Eidechsen Der Dalmatinischen Eilande. Leipzig und Wien: Verlag von Franz Deuticke; 1926.

29. While GM, Michaelides S, Heathcote RJP, et al. Sexual selection drives asymmetric introgression in wall lizards. Ecology Letters. 2015;18(12):1366–1375. 10.1111/ele.12531

30. Miñano MR, While GM, Yang W, et al. Climate Shapes the Geographic Distribution and Introgressive Spread of Color Ornamentation in Common Wall Lizards. The American Naturalist. 2021;198(3):379–393. 10.1086/715186

31. Yang W, While GM, Laakkonen H, et al. Genomic evidence for asymmetric introgression by sexual selection in the common wall lizard. Molecular Ecology. 2018;27(21):4213–4224. 10.1111/mec.14861

32. Feiner N, Yang W, Bunikis I, While GM, Uller T. Adaptive introgression reveals the genetic basis of a sexually selected syndrome in wall lizards. Science Advances. 2024;10(14):eadk9315. 10.1126/sciadv.adk9315

33. Heathcote RJP, While GM, MacGregor HEA, et al. Male behaviour drives assortative reproduction during the initial stage of secondary contact. Journal of Evolutionary Biology. 2016;29(5):1003–1015. 10.1111/jeb.12840

34. MacGregor HEA, While GM, Barrett J, et al. Experimental contact zones reveal causes and targets of sexual selection in hybridizing lizards. Functional Ecology. 2017;31(3):742–752. 10.1111/1365-2435.12767

35. Nagase T, Sanai Y, Nakamura S, Asato H, Harii K, Osumi N. Roles of HNK-1 carbohydrate epitope and its synthetic glucuronyltransferase genes on migration of rat neural crest cells. Journal of Anatomy. 2003;203(1):77–88. 10.1046/j.1469-7580.2003.00205.x

36. Bronner-Fraser M. Perturbation of cranial neural crest migration by the HNK-1 antibody. Developmental Biology. 1987;123(2):321–331. 10.1016/0012-1606(87)90390-3

37. Clark K, Bender G, Murray BP, et al. Evidence for the neural crest origin of turtle plastron bones. genesis. 2001;31(3):111–117. 10.1002/gene.10012

38. Giovannone D, Ortega B, Reyes M, et al. Chicken trunk neural crest migration visualized with HNK1. Acta Histochemica. 2015;117(3):255–266. 10.1016/j.acthis.2015.03.002

39. Hirata M, Ito K, Tsuneki K. Migration and Colonization Patterns of HNK-1-Immunoreactive Neural Crest Cells in Lamprey and Swordtail Embryos. jzoo. 1997;14(2):305–312. 10.2108/zsj.14.305

40. Juarez M, Reyes M, Coleman T, et al. Characterization of the trunk neural crest in the bamboo shark, Chiloscyllium punctatum. Journal of Comparative Neurology. 2013;521(14):3303–3320. 10.1002/cne.23351

41. Kalcheim C, Le Douarin NM. Requirement of a neural tube signal for the differentiation of neural crest cells into dorsal root ganglia. Developmental Biology. 1986;116(2):451–466. 10.1016/0012-1606(86)90146-6

42. Olsson L, Moury JD, Carl TF, Håstad O, Hanken J. Cranial neural crest-cell migration in the direct-developing frog, Eleutherodactylus coqui: Molecular heterogeneity within and among migratory streams. Zoology. 2002;105(1):3–13. 10.1078/0944-2006-00051

43. Erickson CA, Loring JF, Lester SM. Migratory pathways of HNK-1-immunoreactive neural crest cells in the rat embryo. Developmental Biology. 1989;134(1):112–118. 10.1016/0012-1606(89)90082-1

44. Tucker GC, Delarue M, Zada S, Boucaut JC, Thiery JP. Expression of the HNK-1/NC-1 epitope in early vertebrate neurogenesis. Cell Tissue Res. 1988;251(2):457–465. 10.1007/BF00215855

45. Abo T, Balch CM. A differentiation antigen of human NK and K cells identified by a monoclonal antibody (HNK-1). The Journal of Immunology. 1981;127(3):1024–1029. 10.4049/jimmunol.127.3.1024

46. Simões-Costa M, Bronner ME. Establishing neural crest identity: a gene regulatory recipe. Development (Cambridge, England). 2015;142(2):242–257. 10.1242/dev.105445

47. Qin Q, Xu Y, He T, Qin C, Xu J. Normal and disease-related biological functions of Twist1 and underlying molecular mechanisms. Cell Res. 2012;22(1):90–106. 10.1038/cr.2011.144

48. Soldatov R, Kaucka M, Kastriti ME, et al. Spatiotemporal structure of cell fate decisions in murine neural crest. Science. 2019;364(6444):eaas9536. 10.1126/science.aas9536

49. Howard AG, Baker PA, Ibarra-García-Padilla R, et al. An atlas of neural crest lineages along the posterior developing zebrafish at single-cell resolution. eLife. 2021;10:1–31. 10.7554/eLife.60005

50. Hanemaaijer ES, Margaritis T, Sanders K, et al. Single-cell atlas of developing murine adrenal gland reveals relation of Schwann cell precursor signature to neuroblastoma phenotype. PNAS. 2021;118(5). 10.1073/pnas.2022350118

51. Kuratani SC, Kirby ML. Initial migration and distribution of the cardiac neural crest in the avian embryo: An introduction to the concept of the circumpharyngeal crest. American Journal of Anatomy. 1991;191(3):215–227. 10.1002/aja.1001910302

52. Kvasilova A, Gregorovicova M, Kundrat M, Sedmera D. HNK-1 in Morphological Study of Development of the Cardiac Conduction System in Selected Groups of Sauropsida. The Anatomical Record. 2019;302(1):69–82. 10.1002/ar.23925

53. Nakagawa M, Thompson RP, Terracio L, Borg TK. Developmental anatomy of HNK-1 immunoreactivity in the embryonic rat heart: co-distribution with early conduction tissue. Anat Embryol. 1993;187(5):445–460. 10.1007/BF00174420

54. Chuck ET, Watanabe M. Differential expression of PSA-NCAM and HNK-1 epitopes in the developing cardiac conduction system of the chick. Developmental Dynamics. 1997;209(2):182–195. 10.1002/(SICI)1097-0177(199706)209:2<182::AID-AJA4>3.0.CO;2-E

55. Ikeda T, Iwasaki K, Shimokawa I, Sakai H, Ito H, Matsuo T. Leu-7 immunoreactivity in human and rat embryonic hearts, with special reference to the development of the conduction tissue. Anat Embryol. 1990;182(6):553–562. 10.1007/BF00186462

56. Freyer L, Aggarwal V, Morrow BE. Dual embryonic origin of the mammalian otic vesicle forming the inner ear. Development. 2011;138(24):5403–5414. 10.1242/dev.069849

57. Lee VM, Sechrist JW, Luetolf S, Bronner-Fraser M. Both neural crest and placode contribute to the ciliary ganglion and oculomotor nerve. Developmental Biology. 2003;263(2):176–190. 10.1016/j.ydbio.2003.07.004

58. Streeter GL. The development of the cranial and spinal nerves in the occipital region of the human embryo. American Journal of Anatomy. 1905;4(1):83–116. 10.1002/aja.1000040106

59. Goldberg S, Venkatesh A, Martinez J, et al. The development of the trunk neural crest in the turtle Trachemys scripta. Developmental Dynamics. 2020;249(1):125–140. 10.1002/dvdy.119

60. Canning CA, Lee L, Irving C, Mason I, Jones CM. Sustained interactive Wnt and FGF signaling is required to maintain isthmic identity. Developmental Biology. 2007;305(1):276–286. 10.1016/j.ydbio.2007.02.009

61. Nothwang HG, Ebbers L, Schlüter T, Willaredt MA. The emerging framework of mammalian auditory hindbrain development. Cell Tissue Res. 2015;361(1):33–48. 10.1007/s00441-014-2110-7

62. Azambuja AP, Simoes-Costa M. A regulatory sub-circuit downstream of Wnt signaling controls developmental transitions in neural crest formation. PLOS Genetics. 2021;17(1):e1009296. 10.1371/journal.pgen.1009296

63. Yoshida T, Vivatbutsiri P, Morriss-Kay G, Saga Y, Iseki S. Cell lineage in mammalian craniofacial mesenchyme. Mechanisms of Development. 2008;125(9):797–808. 10.1016/j.mod.2008.06.007

64. Chai Y, Jiang X, Ito Y, et al. Fate of the mammalian cranial neural crest during tooth and mandibular morphogenesis. Development. 2000;127(8):1671–1679. 10.1242/dev.127.8.1671

65. Barrallo-Gimeno A, Nieto MA. The Snail genes as inducers of cell movement and survival: implications in development and cancer. Development. 2005;132(14):3151–3161. 10.1242/dev.01907

66. Sefton M, Sanchez S, Nieto MA. Conserved and divergent roles for members of the Snail family of transcription factors in the chick and mouse embryo. Development. 1998;125(16):3111–3121. 10.1242/dev.125.16.3111

67. Barriga EH, Maxwell PH, Reyes AE, Mayor R. The hypoxia factor Hif-1α controls neural crest chemotaxis and epithelial to mesenchymal transition. Journal of Cell Biology. 2013;201(5):759–776. 10.1083/jcb.201212100

68. Vincentz JW, Firulli BA, Lin A, Spicer DB, Howard MJ, Firulli AB. Twist1 Controls a Cell-Specification Switch Governing Cell Fate Decisions within the Cardiac Neural Crest. PLOS Genetics. 2013;9(3):e1003405. 10.1371/journal.pgen.1003405

69. Tavares AT, Izpisuja-Belmonte JC, Rodriguez-Leon J. Developmental expression of chick twist and its regulation during limb patterning. Int J Dev Biol. 2004;45(5-6):707–713. 10.1387/ijdb.11669372

70. Liu JAJ, Wu MH, Yan CH, et al. Phosphorylation of Sox9 is required for neural crest delamination and is regulated downstream of BMP and canonical Wnt signaling. Proceedings of the National Academy of Sciences. 2013;110(8):2882–2887. 10.1073/pnas.1211747110

71. McKeown SJ, Lee VM, Bronner-Fraser M, Newgreen DF, Farlie PG. Sox10 overexpression induces neural crest-like cells from all dorsoventral levels of the neural tube but inhibits differentiation. Developmental Dynamics. 2005;233(2):430–444. 10.1002/dvdy.20341

72. Cheng YC, Cheung M, Abu-Elmagd MM, Orme A, Scotting PJ. Chick Sox10, a transcription factor expressed in both early neural crest cells and central nervous system. Developmental Brain Research. 2000;121(2):233–241. 10.1016/S0165-3806(00)00049-3

73. Kos R, Reedy MV, Johnson RL, Erickson CA. The winged-helix transcription factor FoxD3 is important for establishing the neural crest lineage and repressing melanogenesis in avian embryos. Development. 2001;128(8):1467–1479. 10.1242/dev.128.8.1467

74. Lukoseviciute M, Gavriouchkina D, Williams RM, et al. From Pioneer to Repressor: Bimodal foxd3 Activity Dynamically Remodels Neural Crest Regulatory Landscape In Vivo. Developmental Cell. 2018;47(5):608–628.e6. 10.1016/j.devcel.2018.11.009

75. D’Autréaux F, Coppola E, Hirsch MR, Birchmeier C, Brunet JF. Homeoprotein Phox2b commands a somatic-to-visceral switch in cranial sensory pathways. Proceedings of the National Academy of Sciences. 2011;108(50):20018–20023. 10.1073/pnas.1110416108

76. Furlan A, Dyachuk V, Kastriti ME, et al. Multipotent peripheral glial cells generate neuroendocrine cells of the adrenal medulla. Science. 2017;357(6346):eaal3753. 10.1126/science.aal3753

77. Pattyn A, Morin X, Cremer H, Goridis C, Brunet JF. The homeobox gene Phox2b is essential for the development of autonomic neural crest derivatives. Nature. 1999;399(6734):366-370. 10.1038/20700

78. Patthey C, Clifford H, Haerty W, Ponting CP, Shimeld SM, Begbie J. Identification of molecular signatures specific for distinct cranial sensory ganglia in the developing chick. Neural Development. 2016;11(3). 10.1186/s13064-016-0057-y

79. Pattyn A, Morin X, Cremer H, Goridis C, Brunet JF. Expression and interactions of the two closely related homeobox genes Phox2a and Phox2b during neurogenesis. Development. 1997;124(20):4065–4075. 10.1242/dev.124.20.4065

80. Chan WH, Komada M, Fukushima T, Southard-Smith EM, Anderson CR, Wakefield MJ. RNA-seq of Isolated Chromaffin Cells Highlights the Role of Sex-Linked and Imprinted Genes in Adrenal Medulla Development. Sci Rep. 2019;9(1):3929. 10.1038/s41598-019-40501-0

81. Langley K, Grant NJ. Molecular markers of sympathoadrenal cells. Cell Tissue Res. 1999;298(2):185–206. 10.1007/PL00008810

82. Khannoon T, Alvarado C, De Bellard ME. Early Development of The Trunk Neural Crest Cells in the Egyptian Cobra Naja H. Haje. Published online May 26, 2021. 10.21203/rs.3.rs-547365/v1

83. Kundrát M. HNK-1 immunoreactivity during early morphogenesis of the head region in a nonmodel vertebrate, crocodile embryo. Naturwissenschaften. 2008;95(11):1063–1072. 10.1007/s00114-008-0426-4

84. Atkins N, Jones SM, Guillette LJ. Timing of parturition in two species of viviparous lizard: influences of β-adrenergic stimulation and temperature upon uterine responses to arginine vasotocin (AVT). J Comp Physiol B. 2006;176(8):783–792. 10.1007/s00360-006-0100-0

85. Feiner N, Rago A, While GM, Uller T. Signatures of selection in embryonic transcriptomes of lizards adapting in parallel to cool climate. Evolution. 2018;72(1):67–81. 10.1111/evo.13397

86. Andrade P, Pinho C, Pérez i de Lanuza G, et al. Regulatory changes in pterin and carotenoid genes underlie balanced color polymorphisms in the wall lizard. Proceedings of the National Academy of Sciences. 2019;116(12):5633–5642. 10.1073/pnas.1820320116

87. Feiner N. Evolutionary lability in Hox cluster structure and gene expression in Anolis lizards. Evolution Letters. 2019;3(5):474–484. 10.1002/evl3.131

88. Tamura K, Stecher G, Kumar S. MEGA11: Molecular Evolutionary Genetics Analysis Version 11. Molecular Biology and Evolution. 2021;38(7):3022–3027. 10.1093/molbev/msab120

